# Combining mass spectrometric platforms for lipidome investigation - Application to the characterisation of disruptions in the lipid profile of pig serum upon β-agonist treatment

**DOI:** 10.1101/2020.03.20.997189

**Authors:** Jérémy Marchand, Yann Guitton, Estelle Martineau, Anne-Lise Royer, David Balgoma, Bruno Le Bizec, Patrick Giraudeau, Gaud Dervilly

## Abstract

In the last decade, many mass spectrometric fingerprinting methods dedicated to lipidomics have been proposed: either non-targeted approaches, coupled with annotation methods, or targeted strategies, aiming at specifically monitoring a limited number of substances.

In a general public health perspective and through a strategy combining non-targeted and targeted lipidomics MS-based approaches, this study aims at identifying disrupted patterns in serum lipidome upon growth promoter treatment in pig and evaluating the relative contributions of the three platforms involved.

Pig serum samples collected during an animal experiment involving control and treated animals, whose food had been supplemented with ractopamine, were extracted and characterised using three MS strategies: Non-targeted RP LC-HRMS; the targeted Lipidyzer™ platform (differential ion mobility associated with shotgun lipidomics) and a homemade LC-HRMS triglyceride platform.

The three different platforms showed complementarity insight into lipid characterisation, which, applied to a selected set of samples, enabled highlighting specific lipid profile patterns involving various lipid classes, mainly in relation with cholesterol esters, sphingomyelins, lactosylceramide, phosphatidylcholines and triglycerides.

Thanks to the combination of both non-targeted and targeted MS approaches, the exploration of various compartments of the pig serum lipidome could be performed, including commonly characterised lipids (Lipidyzer™), triglyceride isomers (Triglyceride platform) -whose accurate analysis was considered an analytical challenge, and unique lipid features (non-targeted LC-HRMS). Thanks to their respective characteristics, the complementarity of the three tools could be demonstrated for public health purposes, with enhanced lipidome coverage, level of characterisation and applicability.

## 1 Introduction

Over the last decade, lipidomics has grown as a major field with diverse applications, such as health or plant studies [1, 2]. Yet, the lipidome is recognised as a complex compartment of the metabolome, with a wide variety of constituting molecules ‒the number of distinct cellular lipids being estimated to range between 10,000 and 100,000 [3], featuring different physicochemical properties, with concentrations spanning over 8 orders of magnitude [4]. For the extraction of such compartment, multiple methods are reported in the literature [5–7], the modified Bligh and Dyer method using chloroform, methanol and water, being the most common one [8]. Then, in order to characterise the lipidome, mass spectrometry (MS) is recognised as the technique of choice [9] and an array of approaches and acquisition methods have been developed to cope with associated complexity [10]. First, the use of MS without prior separation of the analytes (so-called shotgun lipidomics) has been explored, mostly using ESI or nano-ESI as the ionisation source [11], in particular on quadrupole time-of-flight mass spectrometers [12] and more recently, on hybrid quadrupole Orbitrap spectrometers [13, 14]. In order to minimise the ion suppression phenomenon that arises from the complexity of the involved biological matrices and improve the detection of less abundant species, methods using MS coupled to separation techniques such as ultra-high-performance liquid chromatography (UHPLC-MS) [2] or even ion mobility [15] have also emerged.

Across these numerous methods, two main categories can be distinguished and referred to as targeted approaches ‒where a limited number of specific lipid classes or species are monitored and quantified– and non-targeted approaches ‒where an open-list of compounds is analysed for subsequent identification using annotation tools [16]. The latter provide rich information on the lipidome, as they theoretically allow the measurement of any detectable lipid signals [17], resulting in thousands of features. However, this requires data cleaning steps to remove noise and redundancies (isotopes, adducts). Moreover, the assignment of these signals remains a challenging step of the workflow. In contrast, targeted approaches are more selective, thus increasing confidence in the results, even if the acquired information is much more limited. Globally, across the diverse methods and strategies, it appears that no single workflow is sufficient for a wide and complete lipidome characterisation. In such context, the combination of non-targeted and targeted approaches from various complementary techniques is expected to provide an optimal strategy [18] that would further allow discovering unexpected biomarker signals.

Because the application of certain (forbidden) veterinary drugs in livestock aims at modifying the animal carcass composition for leaner meat promotion, it may be expected a shift of associated lipid profiles which could be characterised through lipidomics [19]. Investigating such specific patterns is of prime importance in a public health context aiming at ensuring safe food for the consumer on a chemical point of view, since those growth promoter agents are known to be harmful for human health[20]. Therefore, studying lipidome disruptions as a consequence of growth promoter application would allow generating new knowledge on the mechanism of action of these anabolic agents and especially discovering relevant biomarkers for more efficient screening of such practices. Indeed, classical detection relies on analytical approaches targeting the administered residue or its direct metabolites, which may not be sufficient against emerging illegal practices such as low-dose cocktails or designer-drugs [21]. In this context, the recent application of metabolomics approaches to identify specific biomarkers has been successful [22–26], thus providing an efficient alternative to classical screening methods. Later, lipidomics was also shown to be relevant while highlighting the disruption of phosphatidylglycerols (PG), phosphatidylethanolamine (PE), phosphatidylcholine (PC), and phosphatidic acid (PA) in bovine serum upon trenbolone/estradiol administration [27] and the disruption of PE, phosphatidylinositol (PI) and sphingomyelin (SM) in muscle tissues collected on ractopamine (RAC) fed pigs [19]. However, these preliminary results did not allow a thorough characterisation of the lipidome, since the non-targeted methods applied lacked proper data validation or extensive lipid coverage. Food safety related applications could benefit from the combination of multiple analytical platforms, thus joining their respective forces.

Here ‒through a concrete example of the lipidomics study of RAC effects in pig serum– a strategy making use of three MS platforms (namely: non-targeted LC-HRMS, Lipidyzer™ and an in-house platform for triglyceride regioisomers) is presented in order to determine changes in lipidomic profiles. Since they differ in technology (ion mobility, LC, HRMS, MS/MS) and approach (targeted, non-targeted), this combination is expected to provide both enhanced lipid coverage reliability in the obtained results. In comparison with other multi-platforms approaches published by other groups[18], this original strategy aims to further enhance TG analysis, using a dedicated platform for quantifying their regioisomeric composition. Although the experimental conditions and samples were selected in order to be in a favourable context where lipidic disruptions are expected, this study is an analytical assessment and does not aim to provide comprehensive new knowledge about biological pathways involved or ractopamine biomarkers. Further, it does not intend to propose validated and standardized protocols and is rather to be considered as an evaluation of different available platforms aiming at sharing with the scientific community results regarding their complementarity and respective expected added values. The results are to be evaluated from the point of view of the ability of platforms to generate complementary results from a qualitative point of view.

## 2 Materials and methods

### 2.1 Animal experiment

The blood samples used in this study were obtained from a previously described ethically approved experiment [24], specifically designed to evaluate the disruptions induced in pig blood serum metabolite profiles upon RAC administration. Two groups constituted of randomly divided five four-month-old female pigs were involved over four weeks. After a 3-day acclimatisation, animals from the treated group were exposed to RAC hydrochloride (Sigma Aldrich) through a 10 ppm daily dose in pre-weighted feed (corresponding to 0.45 mg/kg bw/day). Dosage for each animal was verified through complete eating of the daily portion. Six blood samples were collected, respectively at Day-3 (D3), Day-9 (D9), Day-16 (D16), Day-18 (D18), Day-23 (D23) and Day-29 (D29) for each individual from both groups: control (individuals P1 to P5) and treated (individuals P6 to P10). The samples were then allowed to clot at room temperature in order to obtain serum samples.

QC samples were prepared by pooling the same amount of all collected and carefully homogenised samples.

All samples were prepared into suitable 100μL aliquots and immediately stored at − 20 °C until analysis.

### 2.2 Analytical platforms

To characterise the lipidome as widely as possible and evaluate the added value of combining multiple tools, three mass spectrometry platforms were involved for the analysis of the samples from the animal experiment, each of them providing a different level of characterisation. A first option was the non-targeted analysis using Reversed-Phase Ultra High Performance Liquid Chromatography coupled to mass spectrometry (RP UHPLC-HRMS), completed by the targeted platform Lipidyzer™ (differential ion mobility associated with shotgun lipidomics), dedicated to the quantitative analysis of lipids from several classes and finally an in-house developed LC-HRMS platform able to quantify the regioisomeric composition of triglycerides (TG).

For each platform a dedicated sample preparation protocol was carried out, as described in Table 1. While the non-targeted approach was applied on all samples, the targeted tools were implemented for samples collected at the beginning and end of the animal experiment, based on the results from the former. Each time, specific parameters and processing were used, as well as quality assurance (QA) and quality control procedures (QC), which are summarised in Table 1. The full details of the sample preparation and analysis procedures can be found in Supplementary Material.

**Table 1.**
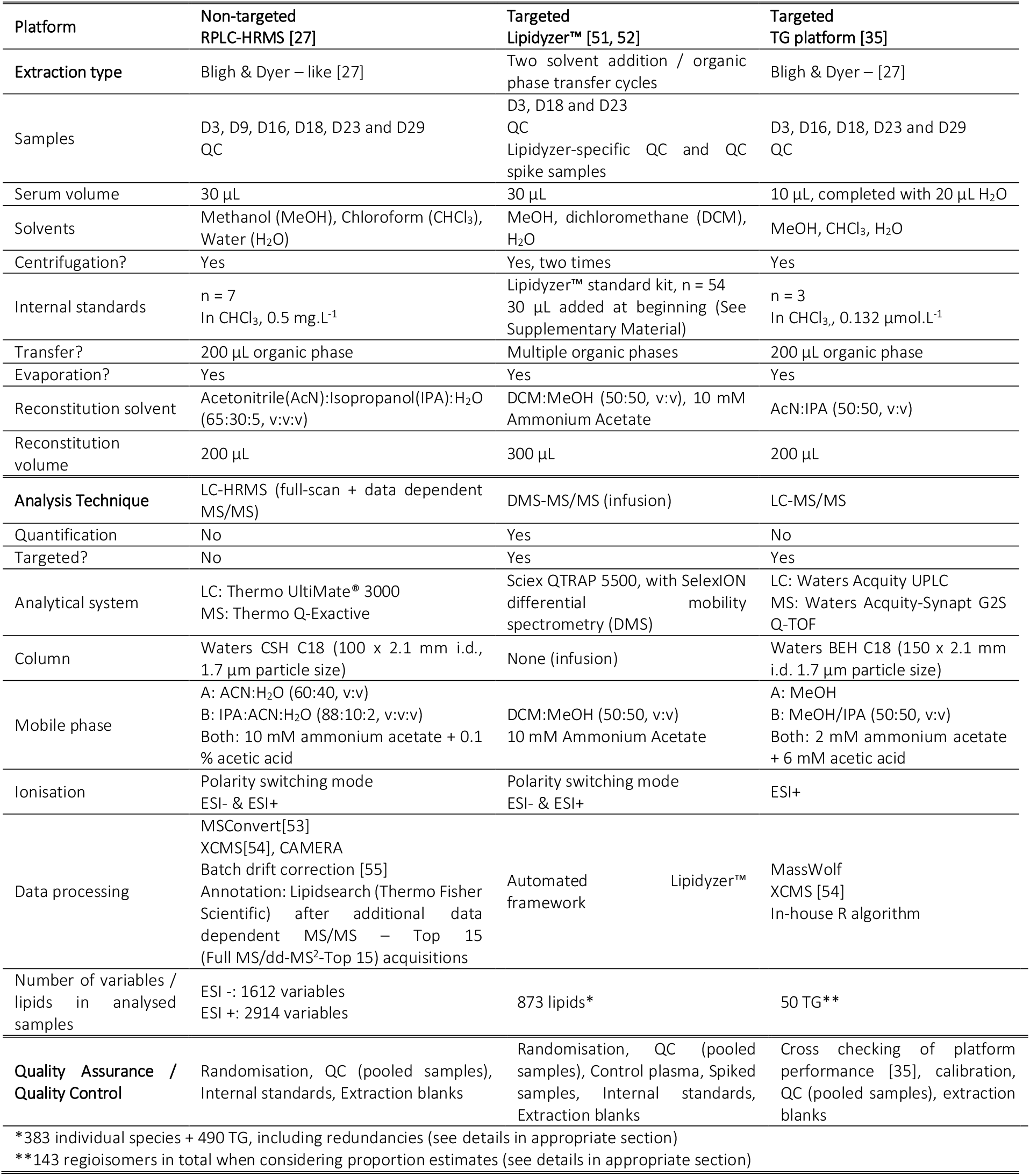
Characteristics of the three used platforms and associated experimental details

| Platform | Non-targeted<br>RPLC-HRMS [27] | Targeted<br>Lipidizer™ [51, 52] | Targeted<br>TG platform [35] |
| --- | --- | --- | --- |
| Extraction type | Bligh & Dyer – like [27] | Two solvent addition / organic phase transfer cycles | Bligh & Dyer – [27] |
| Samples | D3, D9, D16, D18, D23 and D29<br>QC | D3, D18 and D23<br>QC<br>Lipidizer-specific QC and QC spike samples | D3, D16, D18, D23 and D29<br>QC |
| Serum volume | 30 µL | 30 µL | 10 µL, completed with 20 µL H<sub>2</sub>O |
| Solvents | Methanol (MeOH), Chloroform (CHCl<sub>3</sub>), Water (H<sub>2</sub>O) | MeOH, dichloromethane (DCM), H<sub>2</sub>O | MeOH, CHCl<sub>3</sub>, H<sub>2</sub>O |
| Centrifugation? | Yes | Yes, two times | Yes |
| Internal standards | n = 7<br>In CHCl<sub>3</sub>, 0.5 mg.L<sup>-1</sup> | Lipidizer™ standard kit, n = 54<br>30 µL added at beginning (See Supplementary Material) | n = 3<br>In CHCl<sub>3</sub>, 0.132 µmol.L<sup>-1</sup> |
| Transfer? | 200 µL organic phase | Multiple organic phases | 200 µL organic phase |
| Evaporation? | Yes | Yes | Yes |
| Reconstitution solvent | Acetonitrile (ACN):Isopropanol (IPA):H<sub>2</sub>O (65:30:5, v:v:v) | DCM:MeOH (50:50, v:v), 10 mM Ammonium Acetate | AcN:IPA (50:50, v:v) |
| Reconstitution volume | 200 µL | 300 µL | 200 µL |
| Analysis Technique | LC-HRMS (full-scan + data dependent MS/MS) | DMS-MS/MS (infusion) | LC-MS/MS |
| Quantification | No | Yes | No |
| Targeted? | No | Yes | Yes |
| Analytical system | LC: Thermo UltiMate® 3000<br>MS: Thermo Q-Exactive | Sciex QTRAP 5500, with SelexION differential mobility spectrometry (DMS) | LC: Waters Acquity UPLC<br>MS: Waters Acquity-Synapt G2S Q-TOF |
| Column | Waters CSH C18 (100 x 2.1 mm i.d., 1.7 µm particle size) | None (infusion) | Waters BEH C18 (150 x 2.1 mm i.d. 1.7 µm particle size) |
| Mobile phase | A: ACN:H<sub>2</sub>O (60:40, v:v)<br>B: IPA:ACN:H<sub>2</sub>O (88:10:2, v:v:v)<br>Both: 10 mM ammonium acetate + 0.1 % acetic acid | DCM:MeOH (50:50, v:v)<br>10 mM Ammonium Acetate | A: MeOH<br>B: MeOH/IPA (50:50, v:v)<br>Both: 2 mM ammonium acetate + 6 mM acetic acid |
| Ionisation | Polarity switching mode<br>ESI- & ESI+ | Polarity switching mode<br>ESI- & ESI+ | ESI+ |
| Data processing | MSConvert[53]<br>XCMS[54], CAMERA<br>Batch drift correction [55]<br>Annotation: Lipidsearch (Thermo Fisher Scientific) after additional data dependent MS/MS – Top 15 (Full MS/dd-MS<sup>2</sup>-Top 15) acquisitions | Automated framework<br>Lipidizer™ | MassWolf<br>XCMS [54]<br>In-house R algorithm |
| Number of variables / lipids in analysed samples | ESI -: 1612 variables<br>ESI +: 2914 variables | 873 lipids* | 50 TG** |
| Quality Assurance / Quality Control | Randomisation, QC (pooled samples), Internal standards, Extraction blanks | Randomisation, QC (pooled samples), Control plasma, Spiked samples, Internal standards, Extraction blanks | Cross checking of platform performance [35], calibration, QC (pooled samples), extraction blanks |

\*383 individual species + 490 TG, including redundancies (see details in appropriate section)
\*\*143 regioisomers in total when considering proportion estimates (see details in appropriate section)

### 2.3 Data analysis

For non-targeted data, multivariate analysis was performed using SIMCA 13.0.2 (Umetrics AB, Sweden), where log transformation, Pareto scaling and centering were applied. Two-component Principal Component Analyses (PCA) provided an overview of the data and checking the quality of the analysis. Results were then analysed by Partial Least Square Discriminant Analysis (PLS-DA) (centred, UV-scaled). Each PLS-DA model was further validated thanks to permutation tests (n = 100 permutations) and CV-ANOVA. For better interpretability, Orthogonal Projection to Latent Structure Discriminant Analyses (OPLS-DA) were also performed.

Univariate analysis was performed on all datasets using a Wilcoxon test in R studio and *p*-values were calculated using the *coin* package (R studio).

## 3 Results

In order to investigate changes in the lipidome profiles and the complementarity of different MS fingerprinting strategies, a set of samples from which the lipid profiles are expected to be disrupted was chosen as a proof of concept [28]. Below are described and compared the results obtained from three methods: Non-targeted RP UHPLC-HRMS and two targeted approaches, namely Lipidyzer™ and a platform focused on TG regioisomers.

### 3.1 Non-targeted RPLC-HRMS

In the frame of a global lipidomics study, a common method is the non-targeted fingerprinting using LC-HRMS, as it allows studying a large set of lipid species without any *a priori* hypothesis [29] *i.e.* theoretically all lipids accessible to the analysis technique. In the present case, the objective was not to develop a new analytical approach but rather evaluate the contribution of an already established workflow [27] in the frame of a multi-platform study.

After acquisition and verification of the fingerprint quality (see details in Supplementary Material), 1612 and 2914 variables were selected in the ESI− and ESI+ datasets, respectively. A PCA allowed highlighting clustering of the QC samples, thus demonstrating the reproducibility of the analysis (see Supplementary Fig. 1). Furthermore, in PCAs score plots, samples from D3 and D9 did not show major differences between groups, probably because of the slow response of the lipidome to such growth promoting treatment as previously observed [28]. Consequently, these early collection points were removed and the PCAs generated on the resulting ESI+ and ESI− datasets (D16, D18, D23, D29 samples) exhibited separation trends between groups (see Supplementary Fig. 2). PLS-DA were then performed and a discrimination between groups was observed (Fig. 1, left panel) with the following performance: R^2^ = 0.882 and Q^2^ = 0.444 for ESI−; R^2^ = 0.697 and Q^2^ = 0.482 for ESI+. The models were further assessed with CV-ANOVA (*p*-value = 9.5 × 10^−4^ for ESI− and *p*-value = 3.1 × 10^−4^ for ESI+) indicating significant statistical models [30]. For both models, high R^2^ values demonstrated high descriptive ability, while Q^2^ values (< 0.5) pointed out a limited predictability, as confirmed by permutation tests (Supplementary Fig. 3). This was attributed to the high number of variables ‒generating noise‒ while better predictive models are expected through refined selection of variables. Such selection would also answer our needs in terms of classification model practical implementation. Consequently, the variables of interest were determined using a strategy successfully applied in previous works based on OPLS-DA outcomes [23], here through assessment of variable importance for projection of the predictive component (VIPpred) [31], using Workflow4metabolomics 3.3 [32–34]. VIPpred was specifically chosen as it is purely associated with the consequences of the treatment, as opposed to the orthogonal component, associated with the experimental variability and time-related evolution of the individuals. Since a VIPpred > 1.5 threshold returned an excessive number of variables (374 from ESI+; 203 from ESI−), VIPpred > 1.8 was therefore chosen, leading to a selection of 94 and 46 features from the ESI− and ESI+ datasets resp., all of them exhibiting higher signal intensity in the samples from treated animals. From these variables, new PLS-DA models were built (Fig. 1, right panel), showing a strong discrimination between groups, with, the following performance for the reduced ESI+ model: R^2^ = 0.544; Q^2^ = 0.465, CV-ANOVA *p*-value = 5.3 × 10^−4^; and reduced ESI− model: R^2^ = 0.620; Q^2^ = 0.487, CV-ANOVA *p*-value = 3.3 × 10^−4^. The quality of the reduced models was also confirmed by permutation tests (Supplementary Fig. 4).

**Fig. 1.**
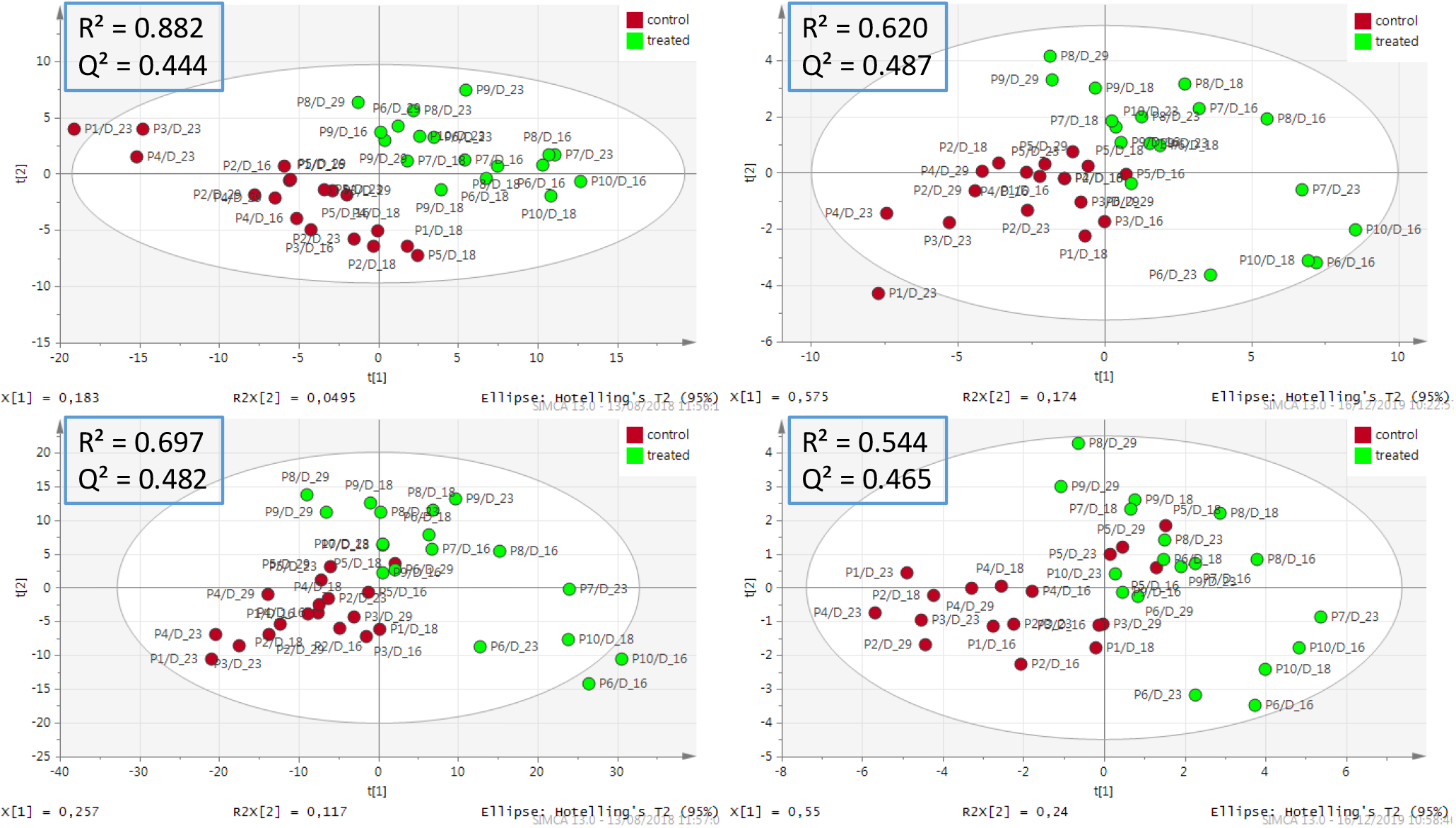
PLS-DA score plots after removing QC, D0, D3, D9 samples from the cleaned ESI− (top) and ESI+ (bottom) datasets acquired with RP UHPLC-HRMS. Left: Datasets containing 1612 (ESI−) and 2914 (ESI+) variables, n = 36. Right: Reduced datasets containing 94 (ESI−) and 46 (ESI+) variables, n = 36. Log 10 transformation, Pareto scaling and centring were applied

The relevance of the selected features was confirmed by “day-by-day” Wilcoxon tests. Thanks to additional data from dependent MS/MS – Top 15 (Full MS/dd-MS^2^-Top 15) experiments performed on QC samples and four typical samples (P4 (control) and P8 (treated) at D18 and D23), a few of them could be putatively using the LipidSearch tool. Annotations and statistical results are detailed in Table 2. Detailed results from LipidSearch for these variables can be found in Supplementary Table 1. From the reduced ESI− dataset, 3 PC, 8 PE and 1 phosphatidylserine (PS) could be annotated whereas 1 PC, 2 PE and 9 TG were annotated from the reduced ESI+ dataset, including 1 PE which was annotated in both ionisation modes (PE(17:0_20:4)). From these preliminary results, it can be noticed that the discrimination between samples from control and treated animals mainly relies on phospholipids and TG, which is consistent with recent literature [19]. PC appear to be mostly discriminant (*p*-value ≤ 0.05) at D16, D18 and D29, PE at D16 and D23 and TG at D23. The annotated phosphatidylserine (PS(18:2_21:0)) was found to be discriminant at all kinetic points between D16 and D29. However, three annotated TG (TG(16:0_17:0_18:1) and the two adducts of TG(18:0_17:0_18:1)) did not exhibit *p*-values ≤ 0.05 and thus could be regarded as modestly involved in the discrimination between groups.

**Table 2.** Putatively annotated variables of interest extracted from the reduced the LC-HRMS datasets, with associated VIPpred values from the OPLS-DA used for variable selection and *p*-values from a Wilcoxon test. **: *p*-value < 0.01; *: *p*-value ≤ 0.05. †: VIPpred values from the OPLS-DA model based on the 1612 (ESI −) and 2914 variables (ESI+) after removal of QC, D0, D3 and D9

| Variable ID | VIPpred<sup>†</sup> | Annotation (LipidSearch) | MS<sup>2</sup> validation (LipidSearch) | *p*-value D16 | *p*-value D18 | *p*-value D23 | *p*-value D29 |
| --- | --- | --- | --- | --- | --- | --- | --- |
| *ESI<sup>-</sup>* |  |  |  |  |  |  |  |
| M791T491 | 1.81 | [PC(18:1_14:0)+CH<sub>3</sub>COO]<sup>-</sup> | ✓ | 0.117 | *0.027 | 0.117 | *0.034 |
| M805T538 | 1.98 | [PC(15:0_18:1)+CH<sub>3</sub>COO]<sup>-</sup> | ✓ | **0.009 | *0.014 | *0.028 | *0.034 |
| M833T633 | 1.84 | [PC(17:0_18:1)+CH<sub>3</sub>COO]<sup>-</sup> | ✓ | *0.028 | *0.014 | 0.076 | *0.034 |
| M715T534 | 1.92 | [PE(16:0_18:2)-H]<sup>-</sup> | ✓ | 0.117 | 0.221 | **0.009 | 0.480 |
| M717T611 | 2.18 | [PE(16:0_18:1)-H]<sup>-</sup> | ✓ | *0.028 | 0.806 | **0.009 | 0.480 |
| M739T518 | 1.97 | [PE(16:0_20:4)-H]<sup>-</sup> | ✓ | *0.047 | 0.086 | *0.016 | 0.480 |
| M745T705 | 2.05 | [PE(18:0_18:1)-H]<sup>-</sup> | ✓ | 0.076 | 0.142 | *0.047 | 0.289 |
| M753T566 | 2.02 | [PE(17:0_20:4)-H]<sup>-</sup> | ✓ | *0.028 | *0.014 | 0.076 | 0.480 |
| M765T524 | 2.17 | [PE(18:1_20:4)-H]<sup>-</sup> | ✓ | 0.076 | 0.142 | *0.016 | 0.157 |
| M723T563 | 1.90 | [PE(16:0p_20:4)-H]<sup>-</sup> | ✓ | **0.009 | 0.221 | **0.009 | 0.077 |
| M751T659 | 1.84 | [PE(16:0p_22:4)-H]<sup>-</sup> | ✓ | **0.009 | 0.327 | *0.028 | 0.157 |
| M829T472 | 1.80 | [PS(18:2_21:0)-H]<sup>-</sup> | ✓ | *0.016 | *0.050 | *0.028 | *0.034 |
| *ESI<sup>+</sup>* |  |  |  |  |  |  |  |
| M777T719 | 1.95 | [PC(16:0_19:0)+H]<sup>+</sup> | X | *0.047 | *0.027 | 0.175 | 0.289 |
| M755T566 | 1.81 | [PE(17:0_20:4)+H]<sup>+</sup> | ✓ | **0.009 | *0.014 | *0.047 | 0.077 |
| M759T836 | 1.84 | [PE(20:0p_18:1)+H]<sup>+</sup> | ✓ | *0.028 | *0.050 | *0.047 | 0.077 |
| M865T1051 | 1.87 | [TG(16:0_17:0_18:1)+NH<sub>4</sub>]<sup>+</sup> | ✓ | 0.117 | 0.086 | 0.076 | 0.157 |
| M879T1059 | 1.92 | [TG(18:0_16:0_18:1)+NH<sub>4</sub>]<sup>+</sup> | ✓ | 0.076 | 0.142 | *0.028 | 0.157 |
| M891T1051 | 1.84 | [TG(17:0_18:1_18:1)+NH<sub>4</sub>]<sup>+</sup> | ✓ | 0.117 | 0.086 | *0.047 | 0.157 |
| M893T1066 | 1.91 | [TG(18:0_17:0_18:1)+NH<sub>4</sub>]<sup>+</sup> | ✓ | 0.076 | 0.086 | 0.076 | 0.289 |
| M898T1065 | 1.84 | [TG(18:0_17:0_18:1)+Na]<sup>+</sup> | ✓ | 0.117 | 0.086 | 0.076 | 0.157 |
| M921T1080 | 2.35 | [TG(18:0_18:1_19:0)+NH<sub>4</sub>]<sup>+</sup> | ✓ | 0.117 | 0.086 | *0.047 | 0.157 |
| M926T1080 | 1.99 | [TG(18:0_18:1_19:0)+Na]<sup>+</sup> | ✓ | *0.047 | 0.086 | 0.076 | 0.157 |
| M919T1066 | 1.88 | [TG(19:1_18:0_18:1)+NH<sub>4</sub>]<sup>+</sup> | ✓ | *0.047 | 0.142 | *0.016 | 0.077 |
| M924T1066 | 1.84 | [TG(19:0_18:1_18:1)+Na]<sup>+</sup> | ✓ | 0.076 | 0.142 | *0.047 | 0.157 |

### 3.2 Lipidyzer™ platform

In order to provide additional insight into lipids involved in the sample group separation observed above, an alternative MS lipidomics approach was applied. Lipidyzer™ is a commercial lipid quantification tool, based on shotgun lipidomics and benefiting from ion mobility, coming with its own workflow and dedicated framework. As it is based on targeted analysis and differs in the separation mode, complementary results from the non-targeted method presented above are expected. Lipidyzer™ was originally designed for human blood serum and plasma studies but its applicability may be tested for other species. However, since it was used here for porcine serum samples, the associated results cannot be considered as absolute concentrations. Because of the lack of validation for pig samples, the Lipidyzer results detailed in this work are therefore considered as “estimated” concentration. Here, this experiment required a limited number of samples, hence samples collected at D3 (as a basis for comparison), D18 and D23 were characterised with Lipidyzer™, as a result of RPLC-HRMS outcomes described above. From the analysed samples, 795 lipid species were actually measured (*i.e.* above limit of quantification in at least one sample), namely: 26 Cholesterol Esters (CE), 10 Ceramides (CER), 7 Dihydroceramides (DCER), 11 Hexosyl ceramides (HCER), 10 Lactosyl ceramides (LCER), 54 Diacylglycerols (DAG), 26 Free Fatty Acids (FFA), 18 Lysophosphatidylcholine (LPC), 89 PC, 13 Lysophosphatidylethanolamines (LPE), 107 PE, 12 SM and 490 TG. Univariate statistical tests (Wilcoxon, day by day) were performed, showing significant shifts upon RAC treatment for 22 CE, 1 CER, 11 DAG, 1 DCER, 1 FFA, 3 HCER, 3 LCER, 1 LPE, 26 PC, 12 PE, 5 SM and 152 TG (see Table 3). Details about these species can be found in Supplementary Table 2.

**Table 3.** Lipid class analysis results from Lipidyzer™, with associated *p*-values from a Wilcoxon test. **: *p*-value ≤ 0.01; *: *p*-value ≤ 0.05

| Lipid class | *p*-value D3 | *p*-value D18 | *p*-value D23 |
| --- | --- | --- | --- |
| CE | 0.55 | *0.03 | 0.10 |
| CER | 0.22 | 0.11 | 0.31 |
| DAG | 0.42 | 0.20 | **0.01 |
| DCER | 0.42 | 1.00 | 0.22 |
| FFA | 1.00 | 0.20 | 0.42 |
| HCER | 0.15 | *0.03 | 0.69 |
| LCER | 0.69 | *0.03 | **0.01 |
| LPC | 0.06 | 0.34 | 0.84 |
| LPE | 0.15 | 0.11 | 0.55 |
| PC | 0.69 | 0.06 | 0.06 |
| PE | 0.84 | *0.03 | **0.01 |
| SM | 0.22 | *0.03 | 1.00 |
| TG | 0.55 | 0.20 | 0.06 |

When looking at the differences of concentration between samples from control and treated animals at D3 for all measured lipids, only 1 HCER (HCER(24:1)) and 1 PE (PE(O-18:0_18:1)) were shown as significant (*p*-value ≤ 0.05), while 1 PC (PC(16:0_18:0)) and 1 TG (TG42:1-FA14:0) were marginally significantly affected (*p*-value ≤ 0.06). This correlates non-targeted results, where no significant patterns could be observed so early in the experiment. Interestingly, CE, CER, DCER, HCER, LCER, LPE and SM species appeared as significant in the discrimination almost exclusively at D18, while DAG exhibited a significant shift in lipid profiles mainly at D23. The significant shift of species from other classes is distributed evenly between D18 and D23. All the lipids were observed to be more concentrated in the serum of treated animals, except for 1 HCER measured in lower concentration in the serum of treated animals at D3 (HCER(24:1)). Globally, when the number of significant lipid species in either D3, D18 or D23 samples was proportionated to the number of analysed species per class, the most altered classes were CE, (85% of analysed species deemed as significant), SM, (42%), TAG (31%), LCER (30%) PC (29%) and HCER (27%). Examples of boxplots illustrating differences in concentration levels between samples from control and treated groups for 4 particular species are presented in Fig. 2.

**Fig. 2.**
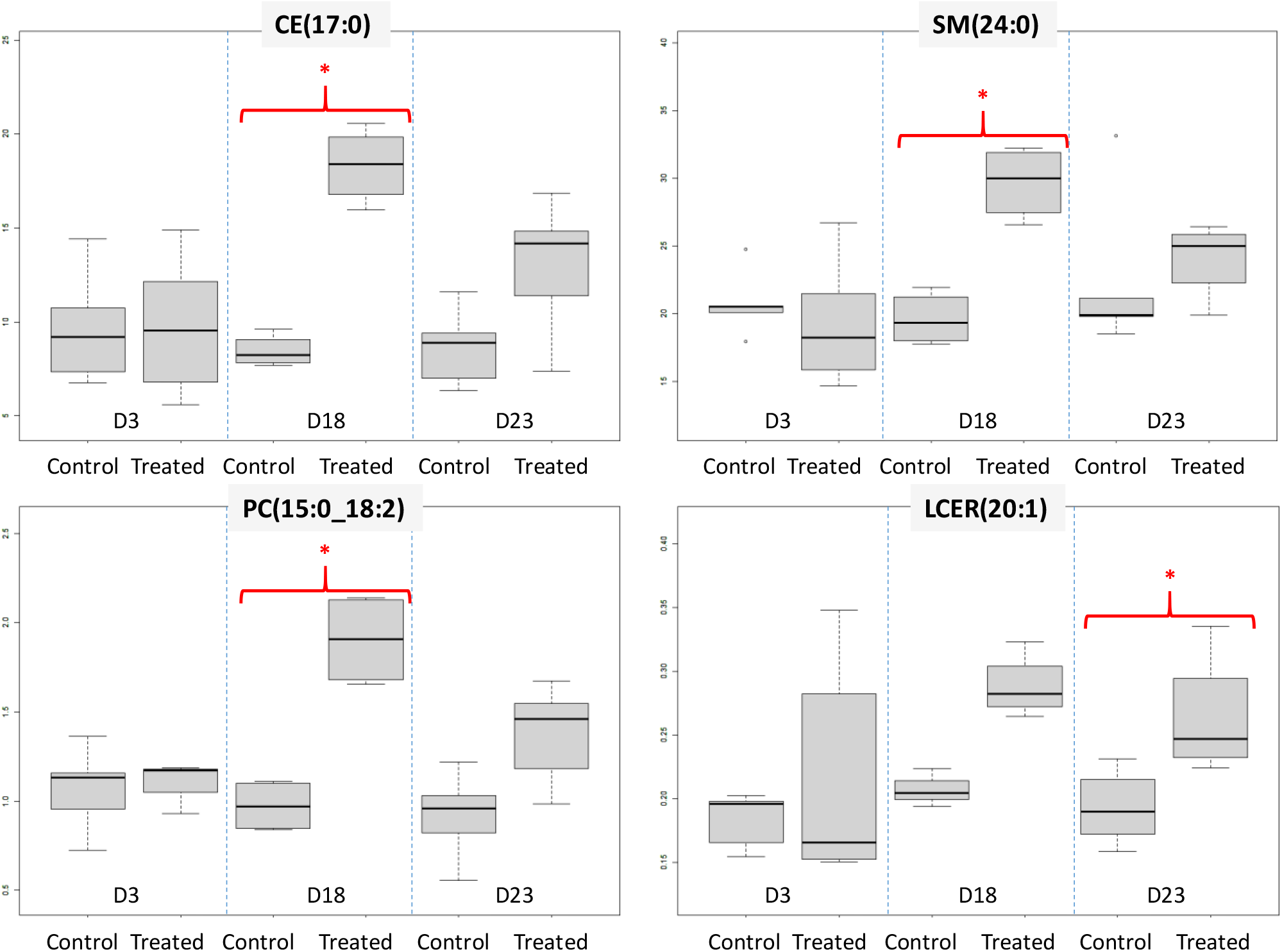
Comparison of estimated concentration (nmol.g^−1^) from four lipid species analysed with Lipidyzer™ between the two animal groups of interest, and for different serum collection points. Here, the quantification cannot be considered as accurate (hence “estimated”) since it is has not been validated on pig serum, as opposed to human. *: *p*-value ≤ 0.05

### 3.3 TG platform

The characterisation of the different TG isomers is an issue which is not completely addressed by Lipidyzer™, which justified resorting to a dedicated TG platform, originally developed for the annotation and semi-quantification of TG isomers in vegetable oils [35].

Through modelling of the fragmentation patterns in TG containing common fatty acids, using multivariate constrained regression, this TG platform is able to determine their regioisomeric composition. This analytical method is semi-quantitative and aims at highlighting TG patterns, together with their fatty acid composition. Relative proportions for each regioisomer (TG(rac-A/B/C); A, B and corresponding to the constituting fatty acyl chains) can also be determined. The analysis was performed on a limited number of relevant samples: D3 as a reference and samples from D16 to D29, corresponding to time points for which most important TG shifts had been observed using both previous platforms.

From univariate statistical tests (Table 4), five TG (TG(52:5), two TG(54:6), TG(54:5) and TG(54:7)) were detected as significant (Wilcoxon test) in the context of the study for the discrimination between control and treated sample groups at D23, with higher concentrations upon RAC treatment. Two of them, namely TG(54:6) at the retention time (Rt) 555.9 s and TG(54:7), were also found as significant at D16, but with a limit *p*-value (0.05) and slightly lower concentrations in treated individuals. For detected TGs, the proportions of the corresponding regioisomers can also be estimated. For instance, the significant variable TG(54:7), detected at Rt 476.03 s was mainly constituted by TG(rac-18:2/18:2/18:3) (around 60%) but also TG(rac-18:2/18:3/18:2) (around 40%).

**Table 4.** Results from the TG platform, with associated *p*-values from a Wilcoxon test. *: *p*-value ≤ 0.05. For each TG signal, the corresponding regioisomers and associated estimated proportions are detailed. The main regioisomers are in bold

| TG_Rt | Corresponding Regioisomers with estimated proportions | *p*-values |  |  |  |  |
| --- | --- | --- | --- | --- | --- | --- |
|  |  | D3 | D16 | D18 | D23 | D29 |
| TG(52:5)_553.44s | TG(rac-18:3/16:0/18:2) ~15%<br>**TG(rac-16:0/18:2/18:3) ~50%**<br>TG(rac-16:0/18:3/18:2) ~35% | 0.44 | 0.77 | 0.64 | *0.03 | 0.06 |
| TG(54:6)_555.9s | **TG(18:2/18:2/18:2)** | 0.17 | *0.05 | 0.39 | *0.03 | 0.72 |
| TG(54:6)_566.5s | **TG(rac-18:3/18:1/18:2) ~60%**<br>TG(rac-18:1/18:2/18:3) ~10%<br>TG(rac-18:1/18:3/18:2) ~30% | 0.17 | 0.18 | 0.25 | *0.03 | 1.00 |
| TG(54:5)_685.8s | **TG(rac-18:3/18:0/18:2) ~60%**<br>TG(rac-18:0/18:2/18:3) ~20%<br>TG(rac-18:0/18:3/18:2) ~20% | 1.00 | 0.65 | 0.15 | *0.05 | 0.51 |
| TG(54:7)_476.03s | **TG(rac-18:2/18:2/18:3) ~60%**<br>TG(rac-18:2/18:3/18:2) ~40% | 0.65 | *0.05 | 0.64 | *0.03 | 1.00 |

## 4 Discussion

### 4.1 Assessment of the complementarity between platforms

Three platforms differently addressing the lipidome were involved in the characterisation of a set of serum samples in which specific lipid patterns are expected to be observed. Although the investigation of pathways involved and ractopamine biomarkers identification was not an objective of the present study, some results may be shared with the scientific community as follows, and rather have to be considered in an analytical perspective. These have to be carefully compared for assessing their respective contributions and complementarity, in lipidomics in general and for the proposed application. As a preliminary step, reproducibility was compared between the platforms, which were assessed by CV(QC)% on common lipid targets (n=30), resulting in median values below 8% which was considered to fit our requirements.

Whatever the platform used, the disruption of various lipid classes could be highlighted in pig serum after several weeks of RAC treatment, as illustrated in Table 5. Same trends could be observed with the three tools, as higher lipid levels were observed in the serum of treated individuals, *e.g.* for TG (non-targeted, Lipidyzer™ and TG platforms) but also for PC and PE (non-targeted and Lipidyzer™ platforms). A graphical illustration of these shared trends can be found in Supplementary Fig. 5.

**Table 5.** Comparison of the results from various MS platform. The analysed lipid classes are mentionned with the level of significance, determined from a univariate Wilcoxon test

|  | Non-targeted RP LC-HRMS |  | Lipidyzer™ |  | TG platform |  |
| --- | --- | --- | --- | --- | --- | --- |
| Class of the relevant Lipids | Analysed & annotated ? | Variation (if significant) | Analysed & annotated ? | Variation (if significant) | Analysed & annotated ? | Variation (if significant) |
| CE | Yes† |  | Yes | ↗D18*<br>↗D23* | No | - |
| CER | Yes† |  | Yes | ↗D18* | No | - |
| DAG | Yes† |  | Yes | ↗D18*<br>↗D23* | No | - |
| DCER | Yes† |  | Yes | ↗D18* | No | - |
| FFA | Yes† |  | Yes | ↗D18*<br>↗D23* | No | - |
| HCER | Yes† |  | Yes | ↘D3*<br>↗D18* | No | - |
| LCER | No |  | Yes | ↗D18*<br>↗D23* | No | - |
| LPC | Yes† |  | Yes | - | No | - |
| LPE | Yes† |  | Yes | ↗D18* | No | - |
| PC | Yes | ↗D16*,<br>↗D18*,<br>↗D23*,<br>↗D29* | Yes | ↗D18*<br>↗D23* | No | - |
| PE | Yes | ↗D16*,<br>↗D18*,<br>↗D23* | Yes | ↗D3*<br>↗D18*<br>↗D23* | No | - |
| PS | Yes | ↗D16*,<br>↗D18*,<br>↗D23*,<br>↗D29* | No | - | No | - |
| SM | Yes† |  | Yes | ↗D18* | No | - |
| TG | Yes | ↗D16*,<br>↗D23* | Yes | ↗D18*<br>↗D23* | Yes | ↘D16*,<br>↗D23* |

Level of significance after Wilcoxon test is indicated with asterisks: \*\*: $p$ -value $\leq 0.01$ ; \*: $p$ -value $\leq 0.05$
†: Lipid class analysed and annotated by non-targeted RP UPLC-HRMS but not observed in the set of selected variables from OPLS-DA (VIPpred > 1.8)
↘: More concentrated in control samples. ↗More concentrated in treated samples. In bold: Days where main disruptions are observed

To check the consistency between these results, the annotated lipids highlighted by the reduced models in non-targeted analysis were searched in Lipidyzer™ outcomes. Most of them could easily be retrieved and were also found to be significant (*p*-value < 0.05) with the same variations towards RAC treatment, highlighting good consistency, in particular for PC(15:0_18:1), PC(17:0_18:1), PC(18:1_14:0) PE(16:0_20:4); PE(16:0_18:2) and PE(16:0p_20:4). The collection dates when these lipids were found to be significant were generally in accordance, although minor differences were observed. For instance, PC(18:1_14:0) was only highlighted at D18 with the non-targeted analysis, whereas it was also found to be marginally significant at D23 (*p*-value = 0.056), using Lipidyzer™ (as “PC(14:0_18:1)”). Still, some lipids that were highlighted with the non-targeted approach were not observed as significant with Lipidyzer™, usually due to a corresponding signal below the limit of quantification with the latter, as observed for PE(17:0_20:4). In other cases, the reason of this difference was less clear; *e.g.* PE(16:0_18:1) and PE(16:0p_22:4) which were retained from ESI− non-targeted results were not found as significant with Lipidyzer™. This could be explained by different measurement bias or by erroneous annotation, even if no obvious inconsistency was observed. Conversely, significant Lipidyzer™ variables were curated in the non-targeted datasets. Even though some lipid classes from which the lipid species were deemed as significant by Lipidyzer™ (*p*-value ≤ 0.05) were annotated in the non-targeted dataset, some did not belong to the set of variables selected for the reduced model. Indeed, CE and DAG were detected and annotated in the ESI+ dataset, LPE and FFA were observed in the ESI− datasets whereas CER, DCER and SM were characterised in both.

An important matter to consider when comparing the results between platforms is their relative capability for lipid annotation, which as a consequence directly influences the biological interpretation With the non-targeted strategy proposed, the annotation is only putative (level 2 or 3 of identification) and a small portion (<20% for both datasets) of the original features could be annotated, thus demonstrating the challenge of this step. Using targeted approaches, such issue is less likely to happen as their workflows were optimised to target specific lipids of interest. Implementing Lipidyzer™ and the TG platforms thus enabled confident lipid assignment.

While comparing the platforms outcomes and lipid annotation, a particular case is the one of TG, where the assignment of the fatty acyl chains (*sn*-1(3) versus *sn*-2) is recognised as a serious analytical challenge, leading to multiple dedicated research studies [36–38].

▪ From the non-targeted method, TG were annotated from their three FA chains (e.g. “TG(16:0_17:0_18:1)”), based on the annotation results from LipidSearch after data dependent MS/MS. Although allowing confident assignment, the results of such approach may in some particular cases be considered with caution as illustrated hereafter. Among the selected variables for instance, some lipids (M926T1080 and M921T1080; highlighted in light grey as well as M898T1065 and M893T1066 highlighted in dark grey in Table 2) were annotated as adducts of the same TG. These variables were initially not discarded during the data processing step because of an inconsistency between the adduct annotation between the CAMERA package and LipidSearch. Also, two other variables (M919T1066 and M924T1066; highlighted in blue in Table 2) were annotated as two different TG when they could potentially be two adducts of the same lipid, as they are isomers of TG(55:2).
▪ In Lipidyzer™, TG results were expressed with the shorthand annotation nomenclature (total number of carbons and unsaturations among the three FA chains and the precision on one of them), such as “TG51:1-FA16:0”. While technically correct, this leads to an overestimation of the TG, as previously highlighted in the literature [18]. Moreover, several Lipidyzer™ candidates (*e.g.* TG51:1-FA18:1 and TG51:1-FA16:0) can correspond to a single TG feature in RP LC-HRMS (*e.g.* TG(16:0_17:0_18:1)), and *vice-versa*, thus complicating result comparison.
▪ Because of previous issues in TG assignment, a dedicated platform for the determination of TG regioisomeric composition was used [35]. It is interesting to note that the TGs highlighted with the dedicated tool were not those annotated in non-targeted data. Moreover, after conversion to the corresponding shorthand annotation to allow such comparison, none of them was deemed as significant with Lipidyzer™, which could be due to the overestimation of TG with the latter. Conversely, none of the discriminant TG highlighted within the RPLC-HRMS results were monitored with the TG platform, since it is designed for the analysis of even FA chains TG only. It is interesting to note that this specific platform allowed obtaining confident results on TG and the position (*sn*-1(3) versus *sn*-2) of their constituting FA chains. Thus, it yielded finer results than the combined use of non targeted and Lipidyzer platforms ‒an approach that was already explored by Contrepois *et al.* [18].

Between all evaluated platforms, Lipidyzer™ offered the most detailed analysis for a large number of lipids, providing a large amount of biologically interpretable data. Yet, interpretation issues were observed when considering the TG, because of the overestimated occurrence of this class, whereas the TG platform could bring information on the regioisomers of interest without doubt. However, the latter was designed for this class only and the number of followed species and regioisomers is limited.

Nonetheless, targeted platforms focus on a limited number of lipids, originally selected for a particular application, *i.e.* the human serum/plasma studies for Lipidyzer™ and vegetable oils for the TG platform. Hence, the relevance of the monitored compounds is not guaranteed when applied to a different research question and species of interest are also likely to be overlooked, as opposed to the non-targeted strategy. For instance, applying the latter enabled highlighting PS(18:2_21:0) in ESI− as well as PC(15:0_16:0) and PC(16:0_19:0) in ESI+ as relevant upon RAC treatment.

Regarding practical considerations, Lipidyzer can be performed in an easy manner, thanks to the entirely software-guided workflow, from instrument calibration to processing. Comparatively, the non-targeted platform requires more expertise, in particular for data processing, even though tools are available to make this step more accessible, such as Workflow4Metabolomics (W4M) [32, 33]. Since it is still recent, the TG platform still requires a high level of expertise for using the dedicated in-house R algorithm.

Among the three platforms, Lipidyzer™ can be considered as the quickest since the analysis (two 15-min injections, comparable with the 30-min of the non-targeted method and 18-min of TG platform), is compensated by the assisted data processing, allowing a direct interpretation of the results. Nevertheless, since it entails the purchase of dedicated instrument/software/kits, Lipidyzer™ implies a substantial financial investment, whereas the other two can be adapted to various instruments types, although buying pure standards is still required, for the confirmation of lipid assignment or the calibration of TG regioisomers.

The investigation of the serum lipidome disruptions upon RAC administration to pigs showed the added value of the three tools. Rather than heaving up one particular platform above the others, these results clearly demonstrate how comprehensive lipidome characterisation is a challenging task, requiring several tools for both enhanced lipidic coverage and increased confidence in the observations.

### 4.2 Biological interpretation

Besides technological assessments in a global lipidomics context, the present work relied on a case study investigating RAC effects on pig blood lipids profile. Although a full biological interpretation of the metabolic pathways was out of the scope of this work, it is worth discussing our results in light of the current knowledge regarding the impact of RAC on metabolism.

RAC is a synthetic drug belonging to the β-agonist family, widely used as a growth promoter in several countries, as it has been shown to improve growing performance such as average daily gain [39] in pigs. However, as such, it is banned in the European Union [40, 41] and robust screening methods are required to detect any potential abuse. In such context, metabolomics has been successfully applied for screening β-agonists treatment in bovine, thus highlighting signature of administration, enabling the construction of new robust models based on these biomarkers [23]. That is why RAC effects have been similarly studied in porcine, using non-targeted tissue screening [19] and serum metabolomics [24]. As the lipids are known to be disrupted by the use of this compound [42–44], the lipidome appears as a promising compartment for inspecting the effects of RAC, prompting their study by NMR lipidomics [28] and the currently presented work. The mechanism of action of RAC as a growth promoter is relatively well-known; it stimulates β_2_-adrenergic receptors, linked with the relaxation of smooth muscles. They enhance the synthesis and decrease the degradation of proteins [45]. A reduction of adipose tissues as effect of RAC treatment is commonly reported, through two pathways: reduction of lipogenesis and/or increase in lipolysis, as reported by Ferreira *et al.* [44]. This review concluded on a predominance of the former, as a treatment generally does not induce an increase of serum non-esterified fatty acids (NEFA), which is characteristic of the latter.

As observed above, the blood serum levels of various lipid classes appear to be affected by the RAC treatment, starting on the third week of experiment. The disrupted phospholipid profiles observed in the present study are in accordance with the NMR study [28] and previous observations on muscle where modified diacylglycerophosphoethanolamine and phosphatidylinositol profiles have been associated with RAC administration to pigs [19]. Among the highlighted classes, the disruption of SM is in accordance with reported observations in tissue, associating changes in sphingomyelin profiles with RAC administration to pigs [19]. For all involved lipid classes, a delay in the action of RAC can be observed. Further, a limited effect is observed at D29, although the animals were still exposed to the drug. Such observation could be hypothesised to be linked with a de-sensitisation regarding the RAC treatment, which occurs from 21 to 28 days according to Ferreira *et al.* [44].

These observations could form the basis for a better understanding of the mechanism underlying β-agonist treatment on lipid metabolism. Here, no particular effect of RAC could be observed on the free FA profiles (covered by Lipidyzer™). Hence, even if a deeper biological interpretation is necessary before drawing definitive conclusions, this seems in accordance with Ferreira’s review [44], suggesting an inhibition of lipogenesis as preferential mechanism of effect of RAC, rather than an increase in lipolysis which would have conducted to higher free FAs levels in blood.

## 5 Conclusion

This work describes the combination of 3 different fingerprinting approaches, in order to join their forces for one single study dedicated to food safety: non-targeted RP LC-HRMS, the targeted Lipidyzer™ platform and a dedicated TG platform. As an application model, serum samples from an animal experiment involving a repartition agent of interest were characterised. This combination allowed a fine characterisation of the lipid profiles, showing particular lipid classes and species disruptions in pig blood serum following RAC treatment. Specific benefits could be highlighted from the three described platforms, in terms of lipidome coverage, level of characterisation or applicability. Although these platforms enabled reaching complementary information, further work should be conducted to validate the proposed workflows.

For optimising lipidome characterisation, next refinements of the strategy will be directed towards the improvement of lipid annotation from non-targeted RP UPLC-HRMS. Many tools have been reported in the recent literature such as LOBSTAHS [46] or LipidMatch [47] and their evaluation/implementation would ensure higher confidence in results and facilitated link with other platforms. The selection of relevant variables could also be improved, through the use of sparse methods [48, 49] or the recent *biosigner* algorithm [50], precisely aiming at building reduced models. Moreover, the TG platform could be extended in order to include more lipid species, thus requiring further developments in order to increase its suitability to a wider range of lipidomics applications. Improvements could also be made for the development of a more user-friendly data processing interface, which would make this platform accessible to less-experienced analysts and accelerate the time dedicated for such data handling.

The described investigations are primarily focused on the use of different analytical platforms and the animal experiment can only be considered as preliminary. Hence, regarding the study of the effects of RAC on pig’s lipidome, further work is still necessary to fully understand the biological implications underlined by the presented results. Additional animal experiments could also be performed involving for instance different dosages or individuals with different characteristics, for confirming these outcomes and validate candidate biomarkers. As a public health perspective, it is expected that the outcomes of the present study may serve risk analysis, either at the risk assessment level while proposing new insight on mode of action and associated effects or at the risk management steps, as the basis for an alternative screening method based on lipid biomarkers.

## Author contributions

GDP, BLB and PG designed research; JM performed research; YG, A-LR and DB contributed analytical tools; JM, EM, YG, GDP and PG analysed data; JM, PG and GDP wrote the paper. All authors read and approved the manuscript.

## Funding

This work was supported by Région Pays de la Loire, FRANCE, through the “Recherche-Formation-Innovation: Food 4.2” program [grant LipidoTool]. The authors would like to thank the Corsaire metabolomics core facility (Biogenouest).

Sciex is also acknowledged for providing access to Lipidyzer™ platform.

## Compliance with ethical standards

The animal study was approved by the national Ethical Committee n°6 (Comité n°6 - Ministère de l’Enseignement Supérieur et de la Recherche – Direction Générale pour la Recherche et l’Innovation – Secrétariat « Autorisation de projet » - 1, rue Descartes, 75231 PARIS cedex 5, France) under agreement 2,015,092,516,084,715 / APAFIS 1914 (protocol number CRIP-2015-054). The study was implemented at Centre de Recherche et d’Investigation Préclinique-CRIP-ONIRIS-Plate-forme de chirurgie et animaleries expérimentales, Oniris, Nantes, France, under agreement number F.44-271.

Institutional and national guidelines were followed for the animal experimentation, as mentioned in Cerfa N° 51706#02 and N° 14906*02; in particular ARTICLES R. 214-87 to 214-137 from CODE RURAL ET DE LA PÊCHE MARITIME (French Regulation).

