## Supplemental for "Combining mass spectrometric platforms for lipidome investigation - Application to the characterisation of disruptions in the lipid profile of pig serum upon β-agonist treatment"

-

a.	Laberca, Oniris, INRAE, 44307, Nantes (France).

b.	EBSI Team, Chimie et Interdisciplinarité : Synthèse, Analyse, Modélisation (CEISAM) Université de Nantes, CNRS, UMR 6230, BP 92208, 2 rue de la Houssinière, 44322 Nantes (France).

c.	SpectroMaitrise, CAPACITES SAS, 26 Bd Vincent Gâche, 44200 Nantes (France)

* Corresponding authors:

Gaud Dervilly-Pinel

LABERCA, Oniris – Site de la Chantrerie

Route de Gachet, CS 4450707

44307 Nantes Cedex 3

FRANCE

Patrick Giraudeau

EBSI Team, Chimie et Interdisciplinarité : Synthèse, Analyse, Modélisation (CEISAM) Université de Nantes, CNRS, UMR 62302 rue de la Houssinière,

44300 Nantes

FRANCE

**Table of contents; Supplementary material**

### Precisions on Material and Methods.

#### 1.1 Solvents and chemicals

All solvents and reagents used for sample preparation were of analytical or HPLC grade quality. Acetic acid and ammonium acetate were purchased from Sigma-Aldrich (St. Quentin Fallavier, France). Water (H_2_O) was purchased from VWR (Strasbourg, France). Methanol (MeOH), Chloroform (CHCl_3_), Dichoromethane (DCM) and Acetonitrile (AcN) were obtained from Fisher Scientific (Elancourt, France). Isopropanol (IPA) was purchased from Grosseron (St Herblain, France).

Internal standards for LC-HRMS analysis were all purchased from Avanti Polar Lipids (Alabaster, Alabama, USA), covering free fatty acids (FFA) lysophosphatidylcholine (LPC), PC, ceramide (CER), PE and triglyceride (TG) classes. LPC (15:0), FFA (15:0), PC (15:0/15:0), FFA (23:0), CER (d18:1/17:0), PE (17:0/17:0) and TG (17:0/17:0/17:0) were thus used.

For the Lipidyzer™ analysis, standards kits dedicated to the Lipidyzer™ platform were provided: The internal standards, containing over 50 deuterated lipids with various FA chains from the 13 analysed lipid classes and subclasses (complete list available in below with their concentrations), QC spike standards, system suitability and SelexION tuning mix. A QC control plasma was also provided in the kit.

**Internal standards used for Lipidyzer**

For this platform, various deuterated lipid standards were used for the quantification of the targeted lipids. This mixture included one or several species from each lipid class as follows (data from the certificate of analysis of the internal standards kit):

| Lipid class | Internal Standards | MW (g.mol^-1^) | Actual Concentration (mg.L^-1^) |
| --- | --- | --- | --- |
| CE | dCE(16:0) | 631.62 | 0.13 |
|  | dCE(16:1) | 629.61 | 0.13 |
|  | dCE(18:1) | 657.64 | 0.57 |
|  | dCE(18:2) | 655.62 | 1.43 |
|  | dCE(20:3) | 681.64 | 0.15 |
|  | dCE(20:4) | 679.62 | 0.18 |
|  | dCE(20:5) | 677.61 | 0.18 |
|  | dCE(22:6) | 703.62 | 0.22 |
| DAG | dDAG(16:0/16:0) | 577.56 | 0.004 |
|  | dDAG(16:0/18:0) | 605.59 | 0.005 |
|  | dDAG(16:0/18:1) | 603.57 | 0.006 |
|  | dDAG(16:0/18:2) | 601.56 | 0.005 |
|  | dDAG(16:0/18:3) | 599.54 | 0.00135 |
|  | dDAG(16:0/20:4) | 625.56 | 0.0015 |
|  | dDAG(16:0/20:5) | 623.54 | 0.00145 |
|  | dDAG(16:0/22:6) | 649.56 | 0.0016 |
| CER | dCER(d16:0) | 546.97 | 0.02 |
| DCER | dDCER(16:0) | 548.99 | 0.004 |
| HCER | dHCER(16:0) | 709.11 | 0.03 |
| LCER | dLCER(16:0) | 871.25 | 0.03 |

| Lipid class | Internal Standards | MW (g.mol^-1^) | Actual Concentration (mg.L^-1^) |
| --- | --- | --- | --- |
| SM | dSM(16:0) | 709.61 | 0.1 |
|  | dSM(18:1) | 735.62 | 0.1 |
|  | dSM(24:0) | 821.73 | 0.1 |
|  | dSM(24:1) | 819.72 | 0.1 |
| FFA | dFFA(16:0) | 265.48 | 0.05 |
|  | dFFA(17:1) | 268.48 | 0.05 |
| LPC | dLPC(16:0) | 504.69 | 0.1 |
| LPE | dLPE(18:0) | 486.64 | 0.05 |
| PC | dPC(16:0/16:1) | 740.6 | 0.0575 |
|  | dPC(16:0/18:1) | 768.63 | 0.2525 |
|  | dPC(16:0/18:2) | 766.62 | 0.255 |
|  | dPC(16:0/18:3) | 764.6 | 0.065 |
|  | dPC(16:0/20:3) | 792.63 | 0.0725 |
|  | dPC(16:0/20:4) | 790.62 | 0.2775 |
|  | dPC(16:0/20:5) | 788.6 | 0.07 |
|  | dPC(16:0/22:4) | 818.65 | 0.075 |
|  | dPC(16:0/22:5) | 816.63 | 0.0775 |
|  | dPC(16:0/22:6) | 814.62 | 0.145 |
| PE | dPE(18:0/18:1) | 750.59 | 0.01 |
|  | dPE(18:0/18:2) | 748.58 | 0.01 |
|  | dPE(18:0/18:3) | 746.56 | 0.0021 |
|  | dPE(18:0/20:3) | 774.59 | 0.0027 |
|  | dPE(18:0/20:4) | 772.58 | 0.01 |
|  | dPE(18:0/20:5) | 770.56 | 0.0022 |
|  | dPE(18:0/22:5) | 798.6 | 0.0024 |
|  | dPE(18:0/22:6) | 796.58 | 0.01 |
| TG | dTG50:1-FA16:0 | 841.81 | 0.13 |
|  | dTG52:1-FA18:0 | 869.84 | 0.14 |
|  | dTG52:2-FA18:1 | 867.82 | 0.14 |
|  | dTG52:3-FA18:2 | 865.8 | 0.14 |
|  | dTG52:4-FA18:3 | 863.79 | 0.04 |
|  | dTG54:4-FA20:3 | 891.82 | 0.04 |
|  | dTG54:5-FA20:4 | 889.8 | 0.04 |
|  | dTG56:7-FA22:6 | 913.8 | 0.038 |

For the Triglyceride platform, various triglyceride standards containing common fatty acids were used: TG(rac-P/O/P), TG(rac-P/P/O), TG(rac-P/L/P), TG(rac-S/P/O), TG(rac-O/P/O), TG(rac-O/O/P), TG(rac-P/L/S), TG(rac-P/O/L), TG(rac-L/L/P), TG(rac-M/O/B), TG(rac-O/L/O), TG(rac-O/O/L), TG(rac-L/S/L), TG(rac-L/O/L), TG(rac-L/L/O), TG(rac-O/O/ALA), and TG(rac-L/L/GLA) were purchased from Larodan (Solna, Sweden), containing lauric acid (La), myristic acid (M), palmitic acid (P), stearic acid (S), arachidic acid (Ar), behenic acid (B), myristoleic acid (ME), palmitoleic acid (PE), oleic acid (O), linoleic acid (L), α-linoleic acid (ALA), γ-linolenic acid (GLA), arachidonic acid (AA), eicosapentaenoic acid (EPA) and docosahexaenoic acid (DHA).

#### 1.2 Sample preparation

##### 1.2.1 Non-targeted workflow

Blood serum samples (D3, D9, D16, D18, D23 and D29) were allowed to thaw on ice, then the lipid fraction was extracted according to a previously published protocol [1]. Briefly, a Bligh and Dyer extraction was performed on 30 µL of blood serum, using methanol (190 µL), a chloroform solution (380 µL) containing internal standards (n = 7, 0.5 mg.L^-1^) and water (120 µL). After centrifugation, 200 µL of the lower phase containing the lipids were transferred in a HPLC vial and evaporated. The final extracts were reconstituted in 200 µL AcN:IPA:H_2_O (65:30:5, v:v:v) before injection.

Two QC samples were also extracted according to the same process. The QC extracts were then mixed together and re-divided in two, to ensure that they were identical.

##### 1.2.2 Lipidyzer™ - Global targeted workflow

For this workflow, the extraction was performed on samples D3, D18 and D23 according to the standardised Lipidyzer™ protocol, using associated chemical standard kits, but was adapted to the available sample amounts, *i.e.* 30 µL of serum instead of 100 µL as recommended by the supplier.

After a thawing period, the samples were mixed with 0.9 mL water, 2mL methanol 0.9 mL of dichloromethane and vortexed. 30 µL of the internal standard mixture were then added before setting the samples for 30 min at room temperature. 1 mL water and 0.9 mL dichloromethane were added, then the samples were vortexed and centrifuged for 10 min until phase separation. The lower phase was transferred in a new tube and 1.8 mL dichloromethane was added to each of the original samples. Once again, the samples were vortexed and centrifuged for 10 min until phase separation. The bottom layers were added to the previous aliquots. The combined lipidic extracts were evaporated under nitrogen and reconstituted in 300 µL of a 10 mM Ammonium Acetate dichloromethane:methanol (50:50) solution.

Three Lipidyzer-specific QC were also prepared, from 0.1 mL of the provided QC control plasma. The standard mixture volume was therefore adapted (0.1 mL). Three Lipidyzer-specific QC spike samples were prepared in the same way, with an addition of 0.05 mL of a QC spike standard before the extraction.

##### 1.2.3 TG targeted workflow

Samples from D3, D16, D18, D23 and D29 were extracted using the same protocol detailed for the non-targeted workflow, using 10 µL of serum, completed to 30 µL with water. The CHCl_3_ solution used for extraction contained TG(10:0/10:0/10:0), TG(14:0/14:0/14:0) and TG(17:0/17:0/17:0) at 0.132 µmol.L^-1^.

After evaporation, the extracts were reconstituted in 200 µL AcN:IPA 50:50 (v:v) before analysis.

QC samples were also extracted, combined and re-divided into separate vials.

#### 1.3 Non-targeted fingerprinting workflow

##### 1.3.1 Analysis method

- Fingerprinting

The lipid extracts were analysed in a randomised order in two batches using an UltiMate® 3000 series high performance liquid chromatography (HPLC) system (Thermo Fisher Scientific, Bremen, Germany) coupled to a hybrid quadrupole-orbitrap (Q-Exactive) mass spectrometer (Thermo Fisher Scientific, Bremen, Germany) equipped with a heated electrospray (H-ESI II) source operating in polarity switching ion mode, allowing the simultaneous acquisition of ESI- and ESI+ datasets.

A full instrument calibration was performed using MSCAL6 ProteoMassT LTQ/FT-Hybrid ESI NEG and MSCAL5 ProteoMassT LTQ/FT-Hybrid ESI POS (Sigma–Aldrich) solutions for the negative and positive modes, respectively. Xcalibur 2.2 (Thermo Fisher Scientific, San Jose, CA, USA) was used for data acquisition and analysis.

For MS acquisition, the following parameters were applied: spray voltage ±3.5 kV; capillary temperature 350 °C; sheath gas flow rate 55 a.u. (arbitrary units), auxiliary gas flow rate 10 a.u., sweep gas flow rate 0; aux gas temperature 300 °C; S-lens RF level 50. The spectra were acquired as profiles in full scan mode, with the mass range (*m/z*) 150 – 1500 and an associated mass resolving power of 35,000 (FWHM) at *m/z* 200.

The chromatographic separation was achieved with a reversed phase CSH C18 (100 x 2.1 mm i.d., 1.7 µm particle size) column (Waters Corporation, Milford, MA) using ACN:H_2_O (60:40) and IPA:ACN:H_2_O (88:10:2) as solvent A and B, respectively; both containing 10 mM ammonium acetate + 0.1 % acetic acid. The programmed gradient was as follows (%A:%B): 60:40 at 0 min; 50:50 at 2 min; 30:70 at 12 min; 1:99 at 17‑25 min; 60:40 at 26-30 min. The flow-rate was maintained at 300 µL/min during the analysis. The column temperature was set at 45°C and the injection volume was 5 µL.

- Annotation

To enable the annotation of the acquired datasets, supplementary injections were performed on the same instrument for two control samples (P4-D18, P4-D23), two treated samples (P8-D18, P8-D23) and a QC.

The samples were analysed according to identical RP-LC-HRMS conditions described above and the Full scan acquisition mode was completed by an additional data dependent MS/MS – Top 15 (Full MS/dd‑MS^2^-Top 15) mode with an exclusion list according to LipidSearch software recommendations. From the full scan spectrum generated, ions of highest intensity (TopN) were then selected in the subsequent scan for fragmentation, thus providing corresponding MS/MS spectra. An exclusion list was set up, in order to avoid the fragmentation of background noise from the mobile phase. For generating this list, full scan injections of the mobile phase were performed, from which the 100 ions of highest intensity on the whole chromatogram were extracted, in positive and negative modes, separately, hence providing a global picture of the background noise. For these acquisitions, the following parameters were applied for both positive and negative ionisation modes unless specified. In Full-scan, the scan range was set from 250 to 1200 *m/z* with resolution 70.000. For the dd-MS^2^ Top-N, the Top-N parameters was set at 15 with a scan range from 200 to 1200 associated with a 35,000 resolving power (FWHM) at *m/z* 200. The normalised collision energy was stepped at 25, 30, 40% for the negative mode and 25, 30% for the positive mode with an isolation window of 1.0 *m/z*.

##### 1.3.2 Data Pre-processing: assessment of the *m/z* instrumental drift

A pre-processing was performed on the acquired data in order to check the precision of the *m/z* measurement along each injection batch. With XCalibur 3.0.63 (Thermo Fisher Scientific, San Jose, CA, USA), the measured *m/z* for each highest intensity ion of the standards was extracted from each QC analysis along the injections, nevertheless discarding the first five injections at the beginning of the batch.

The acquisition of the lipidomics fingerprint was performed with a high resolving power instrument, in order to obtain high accuracy on the *m/z* measurement and a high resolving power between the acquired lipid features. However, this type of high accuracy *m/z* measurement can be sensitive to many parameters across an analysis batch, resulting in a drift of the measured value, even when a *m/z* calibration is performed before every batch. Therefore, any drift of the *m/z* measurement was assessed using chemical standards.

Here, internal standards corresponding to several lipids from various classes and uncommon FA chains (resulting in their eventual wide spreading along the chromatogram) were added during the sample preparation step in all the samples, including QCs. As these QC samples were analysed regularly for each injection batch (once every five samples), it is then possible to extract the measured *m/z* values of these standards from the QC samples and compare the measured values against the theoretical ones, hence allowing the assessment of the *m/z* measurement accuracy of the instrument during analysis. A high intensity ion was chosen for each lipid standard, in positive and negative modes when possible. Supplementary Figure 6 shows the mass accuracy, expressed in ppm across the first (A) and the second batch (B), on the various QC injections performed during the sequence. Only the last of the first six QC injections at the beginning of each batch were kept, as the first ones are only used to ensure stabilisation of the system [2]. On both injection batches, it can be seen (Supplementary Figure 6) that the *m/z* measurement error is reasonably stable for both modes of ionisation, with very little variation during the multiple QC acquisitions. Thus, the variation does not exceed 2 ppm for the negative mode, as observed for PC (15:0/15:0) in both batches and 1.8 ppm in the positive mode, as observed for TG (17:0/17:0/17:0) in batch 1. However, it can be noted that the measured error appears generally higher in batch 1 (A) compared to batch 2 (B). Indeed, for batch 1, the maximal error reaches almost 6 ppm in positive mode, as observed for CER (d18:1/17:0) and almost 8 ppm in negative mode, as observed for PC (15:0/15:0) whereas for batch 2, the measured error remains below 2.5 ppm for the same compounds in positive (1.3 ppm) and negative (2.2 ppm) mode, respectively. The reasons for such difference in mass accuracy between the two batches are unclear, as the same protocol (ionisation source cleaning, calibration) was applied. Since the samples were randomly distributed in the two batches and that the same processing is subsequently applied on all of them, the maximal error (8 ppm) must be taken into account for all samples. Hence, for the subsequent peak picking step, applied with XCMS, the parameter “mzdiff” was set at 0.008 (= 8 ppm) for all samples, thus meaning that two MS signals observed on two different samples featuring a *m/z* difference under 8 ppm cannot be considered as two distinct features, considering the involved *m/z* measurement error.

##### 1.3.3 Processing of the datasets

The raw files (*.raw) from the analysis were converted to a universal format (*.mzXML) and polarity split (split in positive and negative ionisation mode datafiles) using the MSConvert software [3]. Subsequent processing was performed with the R package XCMS [4], including peak alignment, peak picking, retention time (RT) alignment followed by a peak grouping and integration step. Parameters were applied as follows: CentWave algorithm [5]; peakwidth = 5-35; mzdiff = 0.008; mzwid = 0.008; minfrac = 0.80. The “find isotope” and “find adducts” functions from the CAMERA R package were also used in order to filter XCMS outputs. For the “find adducts” function, an in-house extended adduct list was used. Two datasets were then obtained, one for the negative mode and one for the positive mode.

As the acquisitions were performed in two batches, the inter-batch and within-batch instrumental drift was corrected following the procedure published by van der Kloet *et al.* [6], in order to compensate the induced analytical error.

- Annotation

Later, an automated annotation of the positive and negative Full MS/dd-MS^2^-Top 15 data was carried out with the LipidSearch software (Thermo Fisher Scientific), thanks to *in-silico* database on lipid and lipid fragments. The resulting annotations were then compared to the features detected with XCMS in order to putatively annotate both the positive and negative datasets.

- Cleaning of the datasets

The normalised data matrices (ESI+ and ESI-) containing LC-MS features for all analysed samples were transferred to Excel®. In order to suppress most of the unstable signals, features with coefficients of variation (CV) across the QC > 30% were eliminated. Moreover, features with retention times (t_R_) < 0.7 min (dead volume) or t_R_ > 22 min (column cleaning at the end of each run) were removed.

In order to suppress redundancies within the data matrices, low intensity isotopes and adducts were also discarded. Based on the CAMERA annotations, only the most abundant ion (M) was kept, as opposed the other isotopes (M+1, M+2, M+3…) which were suppressed from the dataset, while verifying the consistency of the isotope annotation with a manual mass difference calculation. When facing several adducts of a same compound, consistently annotated with both CAMERA and LipidSearch, then only one adduct was kept, on the basis of its abundance, stability (lower CV_QC_), in accordance with the literature [7].

Therefore, for the ESI+ dataset the kept adducts were as follows. For DAG and TG: [M+NH4]^+^; LPC, PC and SM: [M+H]^+^; PI: [M+Na]^+^. For ESI- dataset, the kept adducts were as follows: PC, SM, CER: [M-H]^-^. For the other classes only one adduct or none were annotated.

After such data processing and cleaning steps, 1612 and 2914 variables were finally kept in the ESI- and ESI+ datasets, respectively. These data matrices were then studied by multivariate and univariate statistical approaches, in order to characterise the serum lipidome disruptions induced by a ractopamine treatment in pigs.

#### 1.4 Lipidyzer™ platform

- Acquisition

The lipid analysis was performed using the standardised Lipidyzer™ (Sciex, Darmstadt, Germany) platform with its integrated workflow [8, 9] on a Sciex QTRAP 5500 (Sciex, Darmstadt, Germany) equipped with SelexION differential mobility spectrometry (DMS). Two injections in positive/negative switching mode were performed for each sample: a first injection involving ion mobility (SelexION turned on) and a second injection without it (SelexION turned off). Each time, targeted MS/MS experiments were performed. A 10 mM Ammonium Acetate dichloromethane:methanol (50:50, v:v) solution was used for continuous infusion.

- Processing

The processing of the spectra and subsequent calculations were automatically performed by the software. Lipid annotation and quantification were respectively performed through targeted MS/MS analysis and signal comparison between the targeted lipid and internal standard from the same lipid class with close structure (similar FA chain). Result tables were thus generated, featuring lipid species concentrations for all the analysed samples.

The results were also provided as “class concentration”, which are constituted by the addition of all the measured lipid species within a lipid class. Univariate statistical analyses were then performed with R studio.

- Data quality

The injection was performed following the recommendations of the platform. An instrument tuning was first performed with the provided tuning mix, in order to automatically calibrate the optimum compensation voltage in SelexION for each lipid class for achieving their separation. Then, system suitability tests were performed and automatically validated before the analysis of the samples. The provided QC control plasma and QC spike samples were injected at the beginning and the end of the injection batch, which was constituted by injections of randomly order samples extracts and experimental QC every five samples. Upon injection, the QC spike sample and QC control plasma were processed and controlled by the platform, in order to ensure the quality of the subsequently acquired data.

#### 1.5 Triglyceride targeted workflow

For the targeted analysis of TG, a dedicated platform was used, from which the parameters have been described elsewhere [10].

- Acquisition

Lipid extracts were injected (7.5 µL) on an Acquity UPLC System (Waters Corporation, Milford, MA) coupled with an Acquity-Synapt G2S Q-TOF (Waters Corporation, Milford, MA) equipped with an electrospray source, in positive ion mode. Chromatographic separation was achieved on a BEH C18 2.1x150 mm (Waters Corporation, Milford, MA) at 50°C using methanol and methanol/isopropanol 50:50 as solvent A and B respectively, each containing 2 mM ammonium acetate and 6 mM acetic acid. The flow was established at 0.4 mL/min. The gradient started at 100% and changed linearly to 97.5% of A at minute 5, 92.5% at minute 10, 70% at minute 12, 40% at minute 14 and 0.5% from minute 15 to minute 18.

The eluent from the chromatography was subsequently ionised by electrospray in a Synapt G2S Q-TOF. The capillary voltage was set at 3 kV, the source temperature at 120 °C, and sampling cone at 40 V. Desolvation temperature was set at 500 °C and desolvation gas flow at 800 L/h. Separation in the time of flight was carried out in high resolution mode. The adducts detected were [M+NH_4_]^+^ and [M+Na]^+^. The adducts [M+NH_4_]^+^ were fragmented in data-dependent-acquisition mode.

Training of the instrument was performed with triglyceride mixtures, at various concentrations: 0.025; 0.125; 1 and 2 µM respectively.

- Processing

The predictions and calculations were performed with an in-house data processing algorithm, carried out in R 3.3.1 run under RStudio 0.99.902. Briefly, the injection data were converted to mzXML format thanks to the Masswolf software, then loaded into XCMS and processed using in-house tools. The number of carbons and unsaturations of the triacylglycerides were assigned by their mass-to-charge ratio of the adducts [M+NH_4_]^+^ and [M+Na]^+^. The fatty acids that constituted the triacylgycerides were assigned by their neutral losses in the fragmentation experiments. The regioisomeric composition was quantified according the relative intensity of the fragmentation patterns as in Balgoma *et al.* [10].

Univariate statistical analysis was subsequently performed on the resulting dataset.

### Supplementary figures and tables

**Supplementary Fig. 1** PCA Score plot (with QC) from the Non-targeted RP LC-HRMS datasets. Top: ESI-. Bottom: ESI+.

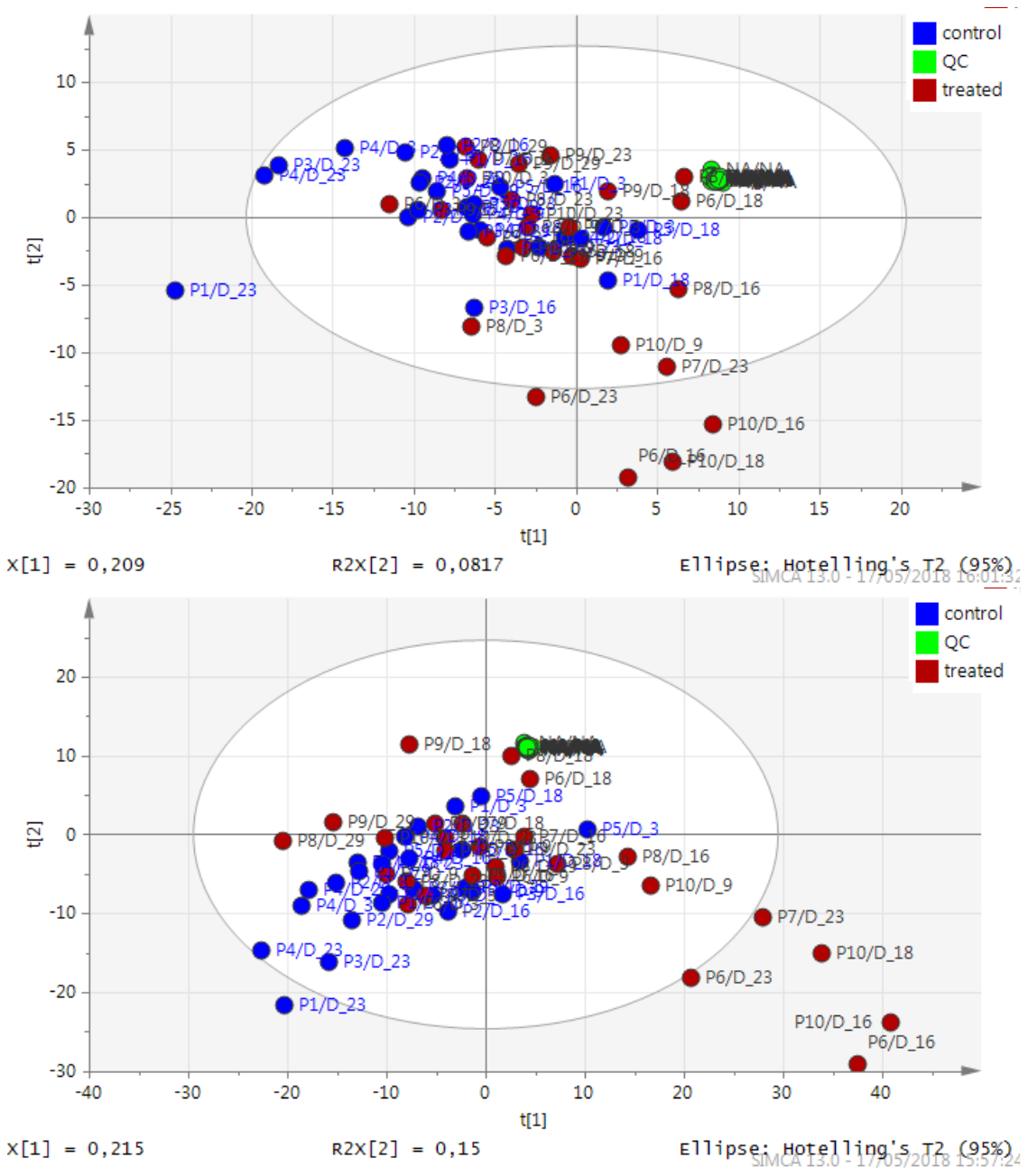

**Supplementary Fig. 2.** PCA Score plot (Without QC, Day¬ 3 and Day-9 samples) from the Non-targeted RP LC-HRMS datasets. Top: ESI-. Bottom: ESI+.

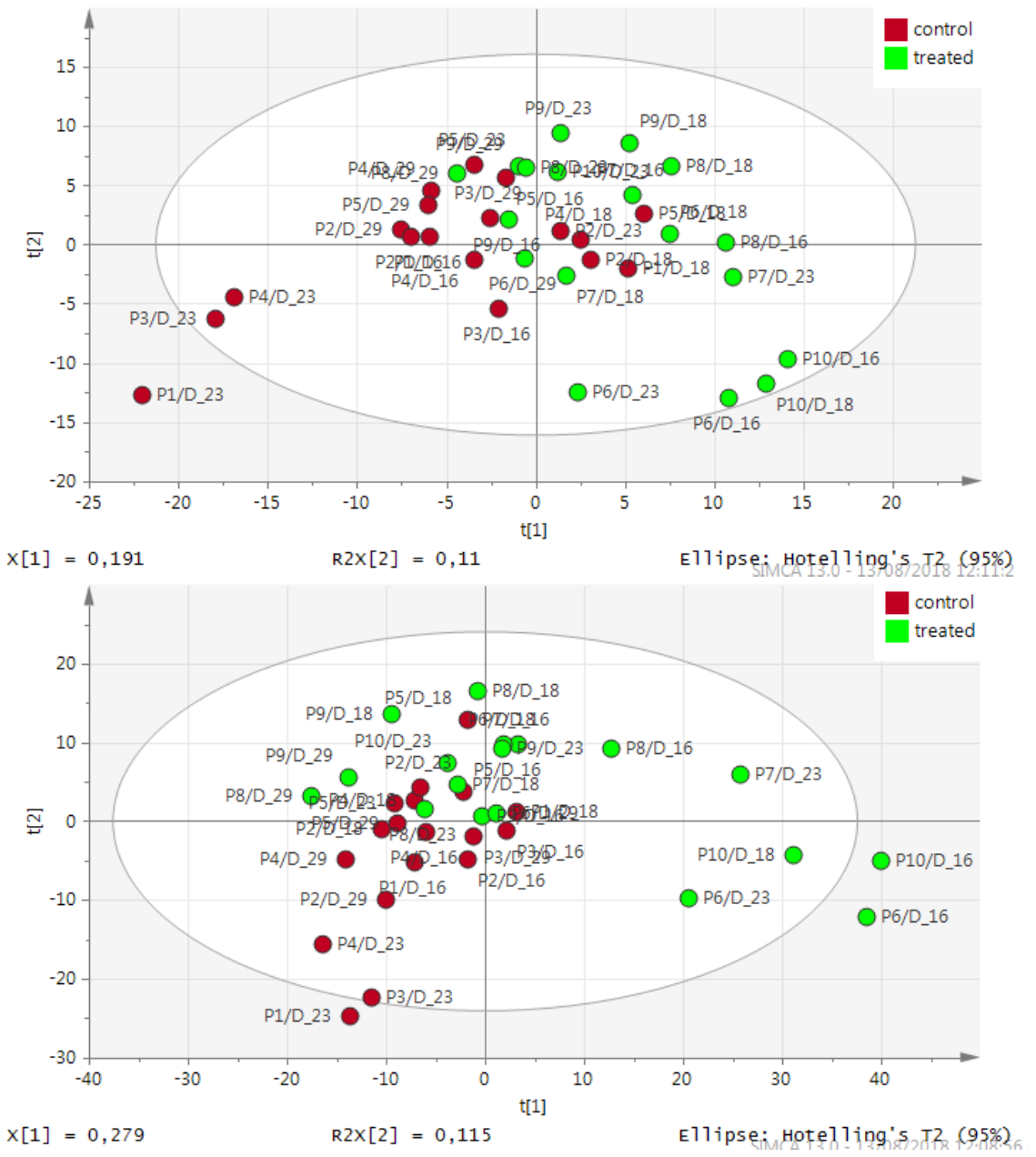

**Supplementary Fig. 3** Permutation tests (n= 100 permutations) associated with the PLS-DA models (Without QC, Day­‑3 and Day-9 samples) from the Non-targeted RP LC-HRMS datasets. Top: ESI-. Bottom: ESI+.

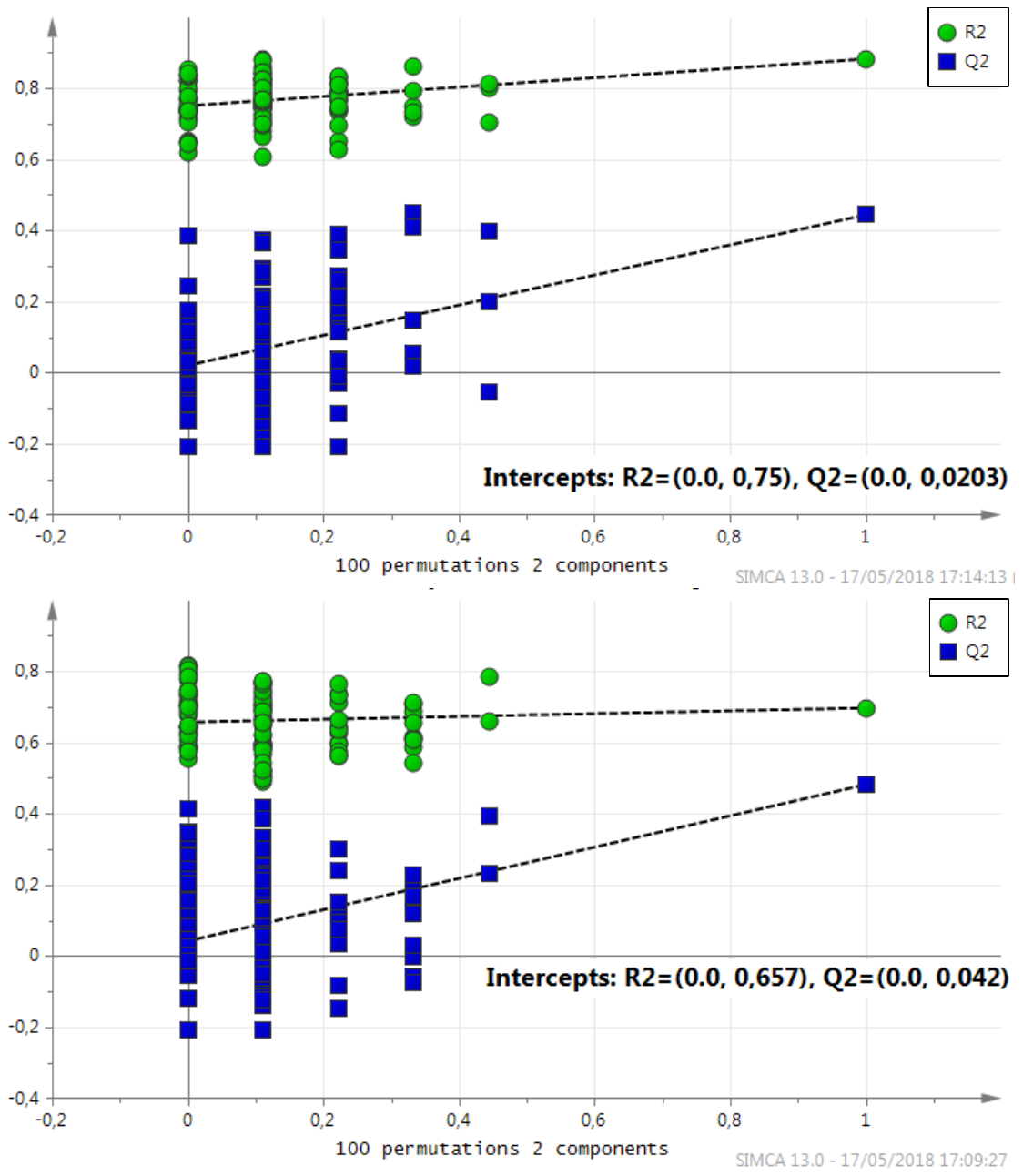

**Supplementary Fig. 4** Permutation tests (n= 100 permutations) associated with the reduced PLS-DA models (Without QC, Day­‑3 and Day-9 samples) from the Non-targeted RP LC-HRMS datasets. Top: ESI-. Bottom: ESI+.

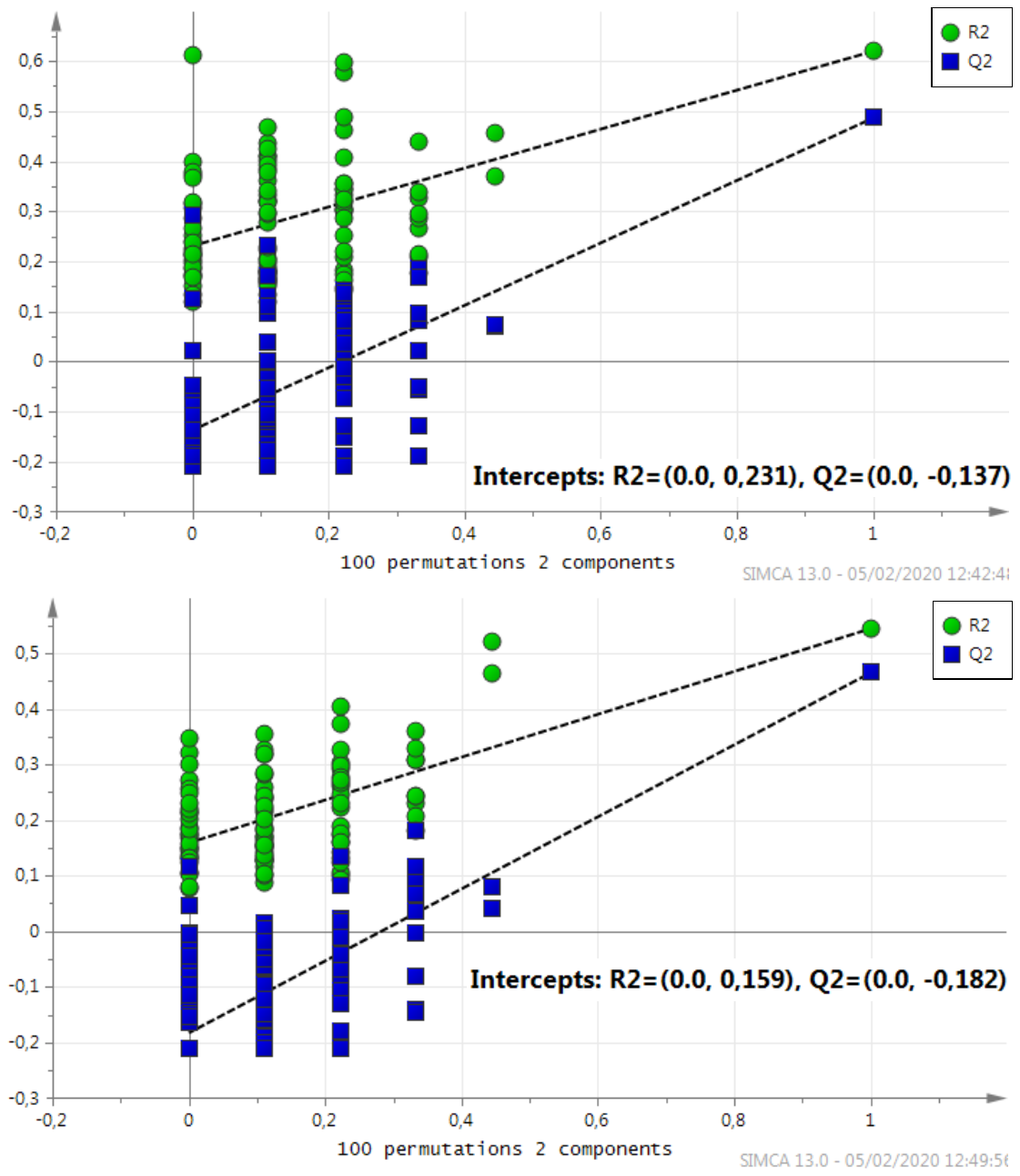

**Supplementary Fig. 5.** Intensity trend comparison of two lipid species analysed with RPLC-HRMS (ESI-) and Lipidyzer in control and treated samples from D23. a.u: arbitrary units

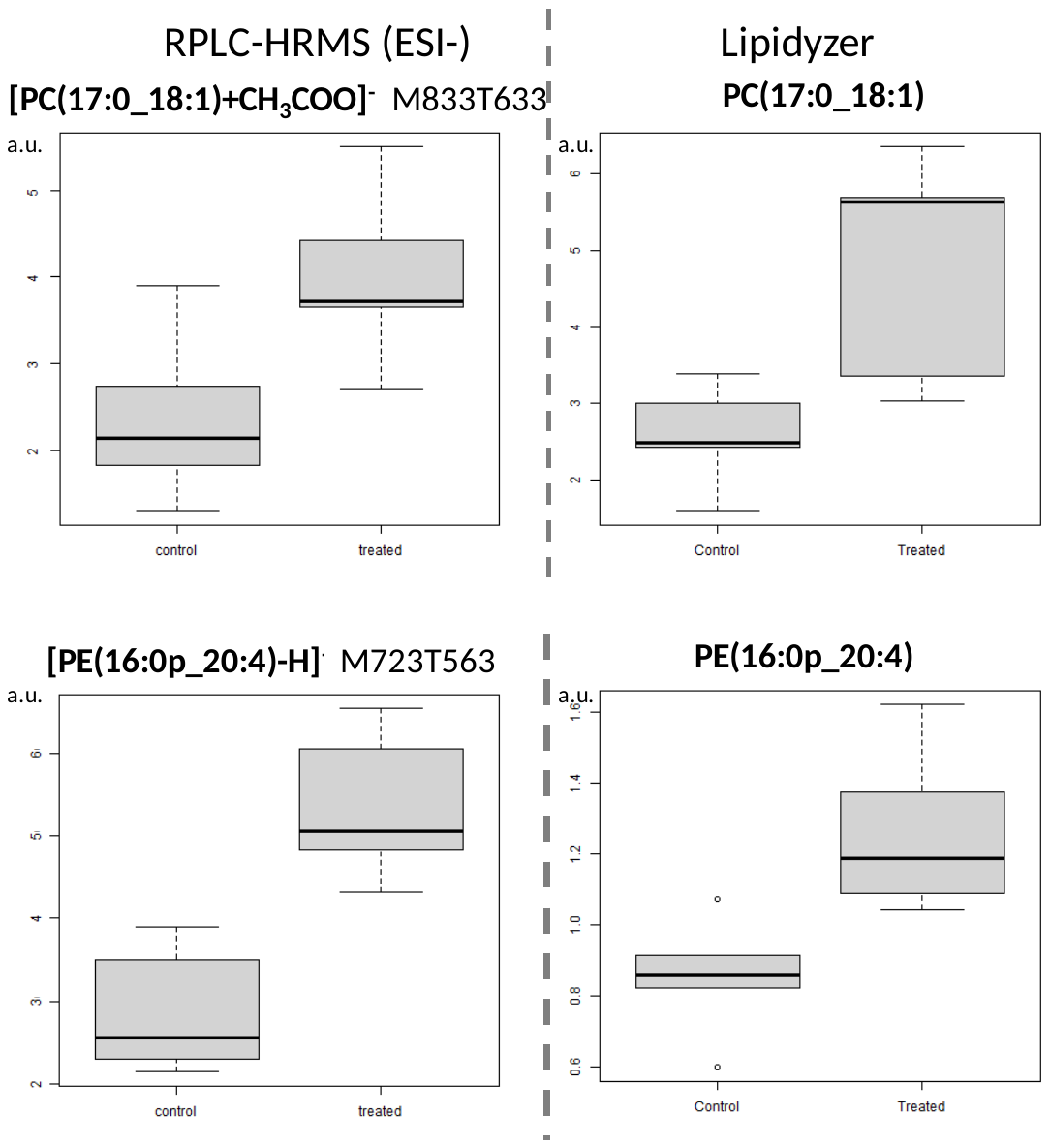

**Supplementary Fig. 6.** m/z measurement error (ppm) on the internal standards signals from QC samples injections. A: Batch 1. B): Batch 2. Solid line: measurements from positive ionisation mode. Dotted line: measurements from negative ionisation mode

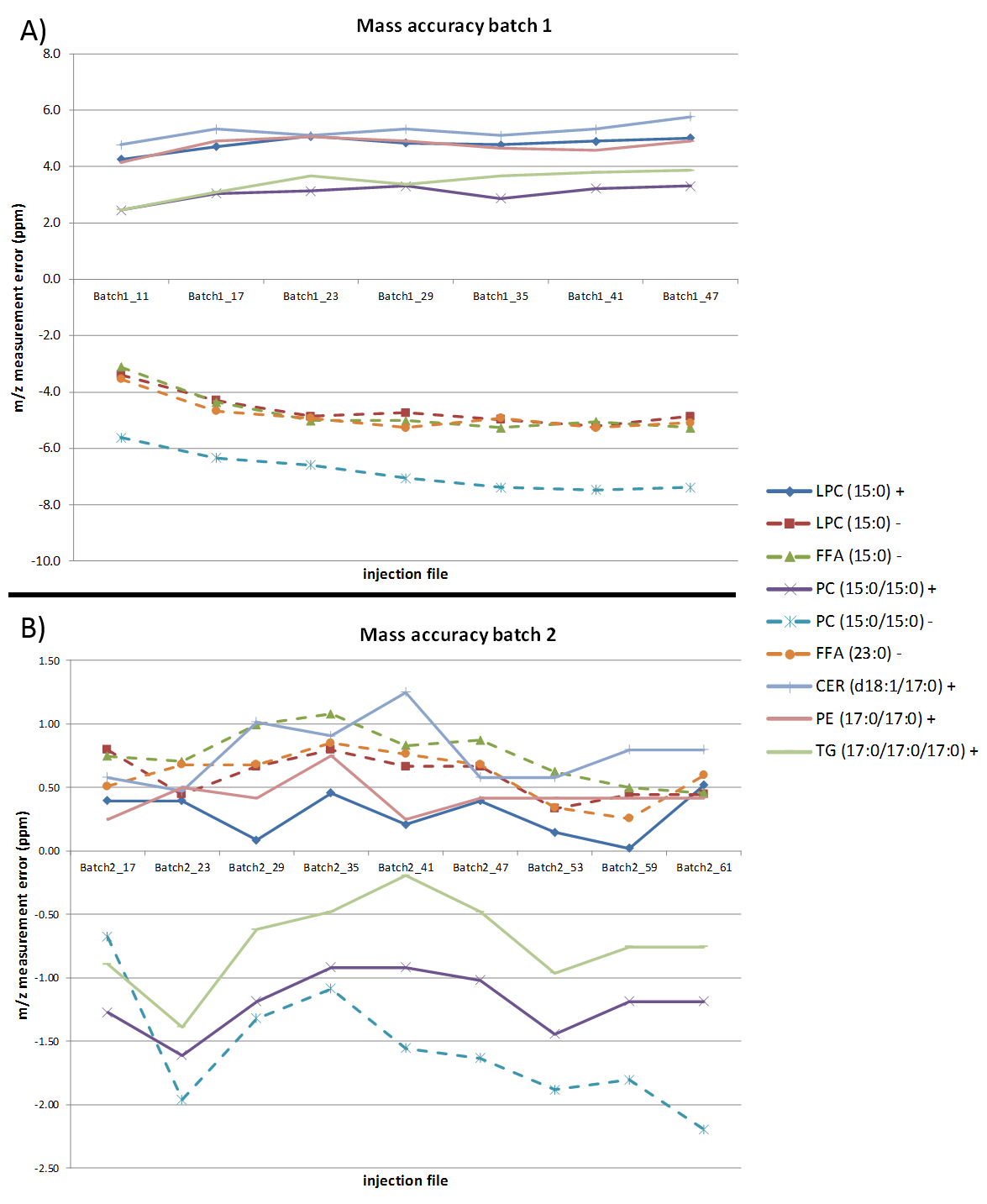

**Supplementary Table 1**. Results from the LipidSearch annotation after MS^2^ analysis for the variables of interest. When possible, annotation was confirmed by both ionisation mode. Bold: variable extracted from the reduced the LC-HRMS datasets; Blue coloring: data from ESI- mode; Red coloring: data from ESI+ mode.

A. *Relevant lipids from* ***ESI****- dataset*

| **Annotation (LipidSearch)** | **Variable ID** | **Observed Mass** | **Calculated Mz** | **Delta (ppm)** | **Product ions** |
| --- | --- | --- | --- | --- | --- |
| **[PC(15:0_18:1)+CH_3_COO]^-^** | M805T538 | 804.57611 | 804.57601053 | 0.8 | (P-Cho)-CH3-H(168.043177):MS2,FA(15:0)-H(241.217422):MS2,FA(18:1)-H(281.248577):MS2,M-CH3(730.540322):MS2 |
| [PC(15:0_18:1)+H]^+^ | M747T538 | 746.56919 | 746.56943347 | -1.1 | C5H12N1(86.096831):MS2,C5H12N(86.096831):MS2,C2H6O4P1(124.999972):MS2,(P-Cho)+H(184.07358):MS2 |
| **[PC(15:0_18:2)+CH_3_COO]^-^** | M803T464 | 802.56070 | 802.560360529999 | 1.1 | (P-Cho)-CH3-H(168.042954):MS2,FA(15:0)-H(241.217373):MS2,FA(18:2)-H(279.232805):MS2,LPC(15:0)-CH3(466.293892):MS2,M-CH3(728.524106):MS2 |
| [PC(15:0_18:2)+H]^+^ | M745T463 | 744.55357 | 744.55378347 | -1.0 | C5H12N1(86.096813):MS2,C5H12N(86.096813):MS2,C2H6O4P1(124.999918):MS2,(P-Cho)+H(184.073515):MS2 |
| **[PC(18:1_14:0)+CH_3_COO]**^-^ | M791T491 | 790.56027 | 790.560360529999 | 0.6 | (P-Cho)-CH3-H(168.043131):MS2,FA(14:0)-H(227.20175):MS2,FA(18:1)-H(281.248483):MS2,LPC(14:0)-CH3(452.278539):MS2,M-CH3(716.524265):MS2 |
| [PC(18:1_14:0)+Na]^+^ | M755T490 | 754.53521 | 754.53572847 | -1.4 | C5H11(71.086078):MS2,C6H9(81.070265):MS2,C5H12N(86.096827):MS2,C7H11(95.085717):MS2,C7H13(97.101547):MS2,C8H13(109.101371):MS2,C2H5O4NaP(146.981935):MS2,M+Na-N(CH3)3-FA(14:0)(467.254354):MS2,NL[PC,+Na]+H(549.488093):MS2,NL[PC](571.469918):MS2,M+Na-N(CH3)3(695.462082):MS2,M+Na(754.534023):MS2 |
| **[PC(17:0_18:1)+CH_3_COO]^-^** | M833T633 | 832.60741 | 832.60731053 | 0.8 | (P-Cho)-CH3-H(168.043005):MS2,FA(17:0)-H(269.248599):MS2,FA(18:1)-H(281.248488):MS2,FA(18:1)-H [isotope](282.251102):MS2,LPC(17:0)-CH3(494.326085):MS2,M-CH3(758.570933):MS2 |
| [PC(17:0_18:1)+Na]^+^ | M797T634 | 796.58266 | 796.58267847 | -0.8 | C6H9(81.070367):MS2,C6H11(83.085982):MS2,C6H13(85.101625):MS2,C5H12N(86.096806):MS2,C7H11(95.085707):MS2,C7H13(97.101395):MS2,C8H13(109.101294):MS2,C9H15(123.116844):MS2,C2H5O4NaP(146.981911):MS2,M+Na-N(CH3)3-FA(17:0)(467.253622):MS2,NL[PC,+Na]+H(591.535375):MS2,NL[PC](613.517367):MS2,M+Na-N(CH3)3(737.509433):MS2,M+Na(796.58364):MS2 |
| **[PE(16:0_18:1)-H]^-^** | M717T611 | 716.52302 | 716.523580529999 | -0.0 | PH(Ethanolamine)-H(140.011827):MS2,M-FA1-FA2+H2O-H(196.037673):MS2,FA(16:0)-H(255.233083):MS2,FA(18:1)-H(281.248391):MS2 |
| [PE(16:0_18:1)+H]^+^ | M719T611 | 718.53657 | 718.53813347 | -2.9 | C5H11(71.086083):MS2,C5H11 [isotope](72.165899):MS2,C6H9(81.070404):MS2,C6H11(83.085966):MS2,C6H13(85.101546):MS2,C7H9(93.070205):MS2,prod(95.0857)(95.085799):MS2,C7H11(95.085799):MS2,C7H13(97.101444):MS2,C8H13(109.101316):MS2,C8H15(111.116828):MS2,C9H13(121.101309):MS2,C9H15(123.116867):MS2,C10H15(135.116928):MS2,C10H17(137.132575):MS2,C11H17(149.132567):MS2,C11H19(151.14864):MS2,NL[PE](577.519183):MS2 |

*(continued)*

| [PE(16:0_18:2)-H]^-^ | M715T534 | 714.50789 | 714.50793053 | 0.7 | PH(Ethanolamine)-H(140.011742):MS2,M-FA1-FA2+H2O-H(196.038408):MS2,FA(16:0)-H(255.232972):MS2,FA(18:2)-H(279.232858):MS2,LPE(16:0)-H(452.279522):MS2 |
| --- | --- | --- | --- | --- | --- |
| [PE(16:0_18:2)+H]^+^ | M717T534 | 716.52373 | 716.52248347 | 1.0 | C6H9(81.070393):MS2,C6H11(83.08592):MS2,C6H13(85.101648):MS2,C6H13 [isotope](86.096824):MS2,C7H9(93.07024):MS2,prod(95.0857)(95.085824):MS2,C7H11(95.085824):MS2,C7H13(97.101351):MS2,C8H11(107.085532):MS2,C8H13(109.101382):MS2,C8H15(111.116857):MS2,C9H13(121.101391):MS2,C9H15(123.117067):MS2,C10H15(135.116804):MS2,MG(16:0)-OH(313.275104):MS2,NL[PE](575.503387):MS2 |
| **[PE(16:0_20:4)-H]^-^** | M739T518 | 738.50763 | 738.50793053 | 0.3 | PH(Ethanolamine)-H(140.011806):MS2,M-FA1-FA2+H2O-H(196.037987):MS2,FA(16:0)-H(255.232915):MS2,FA(20:4)-H-CO2(259.242815):MS2,FA(20:4)-H(303.23285):MS2,LPE(16:0)-H(452.277801):MS2 |
| [PE(16:0_20:4)+H]^+^ | M741T518 | 740.52283 | 740.52248347 | -0.3 | C6H7(79.054695):MS2,C6H9(81.070325):MS2,C6H11(83.086037):MS2,C6H13(85.101598):MS2,C7H9(93.070194):MS2,prod(95.0857)(95.08574):MS2,C7H11(95.08574):MS2,C7H13(97.101364):MS2,C8H11(107.085718):MS2,C8H13(109.101181):MS2,C8H15(111.116913):MS2,C9H13(121.101331):MS2,C9H15(123.116877):MS2,C10H15(135.116753):MS2,C10H17(137.132604):MS2,C11H15(147.116722):MS2,C12H17(161.132541):MS2,C13H19(175.148026):MS2,MG(16:0)-OH(313.273735):MS2,MG(20:4)-OH(361.272479):MS2,NL[PE](599.503667):MS2 |
| **†[PE(17:0_20:4)-H]^-^** | M753T566 | 752.52376 | 752.523580529999 | 1.0 | PH(Ethanolamine)-H(140.01166):MS2,PH(Ethanolamine)-H [isotope](141.016573):MS2,FA(20:4)-H-CO2(259.243253):MS2,FA(17:0)-H(269.248794):MS2,FA(20:4)-H(303.232965):MS2 |
| [PE(17:0_20:4)+H]^+^ | M755T566 | 754.53861 | 754.53813347 | -0.1 | C6H9(81.070514):MS2,C6H13(85.101628):MS2,C7H9(93.070256):MS2,prod(95.0857)(95.085788):MS2,C7H11(95.085788):MS2,C7H13(97.101469):MS2,C7H13 [isotope](98.088545):MS2,C8H11(107.085682):MS2,C8H13(109.101219):MS2,C9H15(123.116947):MS2,C9H15 [isotope](124.027236):MS2,MG(17:0)-OH(327.28999):MS2,NL[PE](613.518993):MS2 |
| **[PE(18:1_20:4)-H]^-^** | M765T524 | 764.52351 | 764.523580529999 | 0.6 | PH(Ethanolamine)-H(140.012087):MS2,FA(20:4)-H-CO2(259.243162):MS2,FA(18:1)-H(281.248471):MS2,FA(20:4)-H(303.232975):MS2 |
| ‡Unconfirmed | | | | | |
| ***[PE(16:0p_20:4)-H]^-^** | M723T563 | 722.51314 | 722.51301553 | 0.9 | PH(Ethanolamine)-H(140.011946):MS2,M-FA1-FA2+H2O-H(196.037847):MS2,FA(20:4)-H-CO2(259.243034):MS2,FA(20:4)-H(303.23301):MS2,LPE(16:0p)-H3O(418.272001):MS2,LPE(16:0p)-H(436.283527):MS2,M-H(722.513054):MS2 |
| ‡Unconfirmed | | | | | |
| **[PE(16:0p_22:4)-H]^-^** | M751T659 | 750.54437 | 750.54431553 | 0.8 | PH(Ethanolamine)-H(140.011806):MS2,M-FA1-FA2+H2O-H(196.037924):MS2,FA(22:4)-H-CO2(287.275014):MS2,FA(22:4)-H(331.264281):MS2,LPE(16:0p)-H3O(418.273626):MS2,LPE(16:0p)-H(436.283523):MS2 |
| ‡Unconfirmed | | | | | |

*(continued)*

| **[PE(18:0_18:1)-H]^-^** | M745T705 | 744.55538 | 744.55488053 | 1.4 | PH(Ethanolamine)-H(140.011877):MS2,PH(Ethanolamine)-H [isotope](141.017073):MS2,M-FA1-FA2+H2O-H(196.038255):MS2,FA(18:1)-H(281.248423):MS2,FA(18:0)-H(283.264011):MS2,LPE(18:0)-H(480.307922):MS2 |
| --- | --- | --- | --- | --- | --- |
| [PE(18:0_18:1)+Na]^+^ | M769T705 | 768.55242 | 768.55137847 | 0.6 | prod(95.0857)(95.085888):MS2,C7H11(95.085888):MS2,H3O4P1Na1(120.966331):MS2,(P-Etr)+Na(164.008335):MS2,NL[PE,+Na]+H(605.54905):MS2,NL[PE](627.528446):MS2 |
| **[PS(18:2_21:0)-H]^-^** | M829T472 | 828.57625 | 828.57601053 | 1.0 | FA(18:2)-H(279.232736):MS2 |
| ‡Unconfirmed | | | | | |
| † Lipid relevant from both ESI+ and ESI- datasets  ‡ Lipid class preferentially ionised in ESI negative mode [11, 12], a lack of confirmation by the positive mode is therefore expected, in particular upon low concentration | | | | | |

B. *Relevant lipids from* ***ESI+*** *dataset*

| **Annotation (LipidSearch)** | **Variable ID** | **Observed Mass** | **Calculated Mz** | **Delta (ppm)** | **Product ions** |
| --- | --- | --- | --- | --- | --- |
| **[PC(16:0_19:0)+H]^+^** | M777T719 | 776.61650 | 776.61638347 | -0.6 | C5H12N1(86.096721):MS2,C5H12N1 [isotope](87.044573):MS2,C5H12N(86.096721):MS2,C5H12N [isotope](87.044573):MS2,(P-Cho)+H(184.073543):MS2 |
| Unconfirmed in ESI- | | | | | |
| **[PE(17:0_20:4)+H]^+^** | M755T566 | 754.53861 | 754.53813347 | -0.1 | C6H9(81.070514):MS2,C6H13(85.101628):MS2,C7H9(93.070256):MS2,prod(95.0857)(95.085788):MS2,C7H11(95.085788):MS2,C7H13(97.101469):MS2,C7H13 [isotope](98.088545):MS2,C8H11(107.085682):MS2,C8H13(109.101219):MS2,C9H15(123.116947):MS2,C9H15 [isotope](124.027236):MS2,MG(17:0)-OH(327.28999):MS2,NL[PE](613.518993):MS2 |
| [PE(17:0_20:4)-H]^-^ | M753T566 | 752.52376 | 752.523580529999 | 1.0 | PH(Ethanolamine)-H(140.01166):MS2,PH(Ethanolamine)-H [isotope](141.016573):MS2,FA(20:4)-H-CO2(259.243253):MS2,FA(17:0)-H(269.248794):MS2,FA(20:4)-H(303.232965):MS2 |
| **[PE(20:0p_18:1)+H]^+^** | M759T836 | 758.60524 | 758.60581847 | -1.5 | C5H9(69.070282):MS2,C6H13(85.101483):MS2,prod(95.0857)(95.085842):MS2,C7H11(95.085842):MS2,C7H13(97.101326):MS2,C8H13(109.101341):MS2,C9H13(121.101214):MS2,MG(18:1)-OH(339.289619):MS2,M-18:1-(CH2=COHCH2OH)(420.324659):MS2 |
| [PE(20:0p_18:1)-H]^-^ | M757T836 | 756.59059 | 756.59126553 | -0.2 | PH(Ethanolamine)-H(140.011762):MS2,FA(18:1)-H(281.248608):MS2,LPE(20:0p)-H3O(474.335218):MS2,LPE(20:0p)-H(492.344371):MS2 |
| **[TG(16:0_17:0_18:1)+NH_4_]^+^** | M865T1051 | 864.80123 | 864.801465469999 | -0.9 | C5H11(71.08607):MS2,C6H9(81.070337):MS2,C6H11(83.085973):MS2,C6H13(85.101574):MS2,C7H9(93.070126):MS2,C7H11(95.085809):MS2,C7H13(97.101418):MS2,C8H11(107.085643):MS2,C8H13(109.101285):MS2,C8H15(111.116958):MS2,C9H13(121.101275):MS2,C9H15(123.116927):MS2,C9H17(125.132399):MS2,C10H15(135.116981):MS2,C10H17(137.132639):MS2,C11H17(149.132666):MS2,C11H19(151.148335):MS2,FA(16:0)-OH(239.237042):MS2,FA(17:0)-OH(253.252824):MS2,FA(18:1)-OH(265.25256):MS2,MG(16:0)-OH(313.273034):MS2,MG(17:0)-OH(327.290695):MS2,MG(18:1)-OH(339.288427):MS2,NL[FA(18:1)-H+NH4](565.518768):MS2,NL[FA(17:0)-H+NH4](577.518282):MS2,NL[FA(16:0)-H+NH4](591.534498):MS2 |
| §Unconfirmed in ESI negative mode | | | | | |
| **[TG(18:0_16:0_18:1)+NH4]^+^** | M879T1059 | 878.81691 | 878.81711547 | -0.9 | C5H7(67.054087):MS2,C5H9(69.069669):MS2,C5H11(71.085275):MS2,C6H7(79.053906):MS2,C6H9(81.069431):MS2,C6H11(83.085029):MS2,C6H13(85.100651):MS2,C7H9(93.069262):MS2,C8H11(107.084761):MS2,C9H15(123.115846):MS2,C11H17(149.131383):MS2,C11H19(151.147873):MS2,FA(16:0)-OH(239.236471):MS2,FA(18:1)-OH(265.251945):MS2,FA(18:0)-OH(267.267794):MS2,MG(16:0)-OH(313.2734):MS2,MG(18:1)-OH(339.289463):MS2,MG(18:0)-OH(341.304933):MS2,NL[FA(18:0)-H+NH4](577.519372):MS2,NL[FA(18:1)-H+NH4](579.535182):MS2,NL[FA(16:0)-H+NH4](605.550954):MS2 |
| §Unconfirmed in ESI negative mode | | | | | |

*(continued)*

| **[TG(18:0_17:0_18:1)+Na]^+^** | M898T1065 | 897.78794 | 897.78816147 | -0.9 | C6H9(81.070436):MS2,C6H11(83.086078):MS2,C6H13(85.101585):MS2,C7H11(95.085821):MS2,C7H13(97.101487):MS2,C8H13(109.101335):MS2,C8H15(111.117136):MS2,C9H13(121.101391):MS2,C9H15(123.116972):MS2,C10H15(135.116874):MS2,C10H17(137.132576):MS2,NL[FA(18:0)-H+Na](591.535345):MS2,NL[FA(18:1)-H+Na](593.551036):MS2,NL[FA(17:0)-H+Na](605.551276):MS2,NL[FA(18:0)](613.517434):MS2,NL[FA(18:1)](615.533218):MS2,NL[FA(17:0)](627.533173):MS2,M+Na(897.788629):MS2 |
| --- | --- | --- | --- | --- | --- |
| §Unconfirmed in ESI negative mode | | | | | |
| **[TG(18:0_17:0_18:1)+NH_4_]^+^** | M893T1066 | 892.83319 | 892.83276547 | -0.1 | C5H11(71.08609):MS2,C6H9(81.070364):MS2,C6H11(83.08596):MS2,C6H13(85.101683):MS2,C7H11(95.085792):MS2,C7H13(97.101476):MS2,C8H13(109.101274):MS2,C8H15(111.116851):MS2,C9H13(121.101208):MS2,C9H15(123.116857):MS2,C10H15(135.117015):MS2,C10H17(137.13271):MS2,FA(18:1)-OH(265.252763):MS2,MG(17:0)-OH(327.288686):MS2,NL[FA(18:0)-H+NH4](591.535349):MS2,NL[FA(18:1)-H+NH4](593.550858):MS2,NL[FA(17:0)-H+NH4](605.550272):MS2 |
| §Unconfirmed in ESI negative mode | | | | | |
| **[TG(17:0_18:1_18:1)+NH_4_]^+^** | M891T1051 | 890.81699 | 890.81711547 | -0.8 | C5H11(71.086077):MS2,C6H7(79.054688):MS2,C6H9(81.070327):MS2,C6H11(83.085983):MS2,C6H13(85.101464):MS2,C7H9(93.07029):MS2,C7H11(95.085783):MS2,C7H13(97.101399):MS2,C8H11(107.08581):MS2,C8H13(109.101278):MS2,C8H15(111.116974):MS2,C9H13(121.101346):MS2,C9H15(123.116956):MS2,C9H17(125.13268):MS2,C10H15(135.116925):MS2,C10H17(137.132683):MS2,C11H17(149.132672):MS2,C11H19(151.148271):MS2,FA(17:0)-OH(253.252386):MS2,FA(18:1)-OH(265.252636):MS2,MG(17:0)-OH(327.288997):MS2,MG(18:1)-OH(339.289382):MS2,NL[FA(18:1)-H+NH4](591.534282):MS2,NL[FA(17:0)-H+NH4](603.534742):MS2 |
| §Unconfirmed in ESI negative mode | | | | | |
| **[TG(18:0_18:1_19:0)+Na]^+^** | M926T1080 | 925.81996 | 925.81946147 | -0.1 | C5H11(71.086014):MS2,C6H9(81.070328):MS2,C6H11(83.086064):MS2,C6H13(85.101596):MS2,C7H11(95.085731):MS2,C7H13(97.10123):MS2,C8H13(109.101056):MS2,C8H15(111.116727):MS2,C9H13(121.101231):MS2,C9H15(123.116775):MS2,NL[FA(19:0)-H+Na](605.549305):MS2,NL[FA(18:0)-H+Na](619.567429):MS2,NL[FA(18:1)-H+Na](621.581921):MS2,NL[FA(19:0)](627.531855):MS2,NL[FA(18:0)](641.54528):MS2,NL[FA(18:1)](643.562477):MS2,M+Na(925.815981):MS2 |
| §Unconfirmed in ESI negative mode | | | | | |
| **[TG(18:0_18:1_19:0)+NH_4_]^+^** | M921T1080 | 920.86438 | 920.864065469999 | -0.3 | C5H9(69.070376):MS2,C5H11(71.086073):MS2,C6H9(81.070296):MS2,C6H11(83.085949):MS2,C6H13(85.101572):MS2,C7H11(95.085782):MS2,C7H13(97.101401):MS2,C8H13(109.101219):MS2,C8H15(111.11674):MS2,C9H13(121.101406):MS2,C9H15(123.116687):MS2,C10H15(135.116719):MS2,NL[FA(19:0)-H+NH4](605.550812):MS2,NL[FA(18:0)-H+NH4](619.566993):MS2,NL[FA(18:1)-H+NH4](621.583058):MS2 |
| §Unconfirmed in ESI negative mode | | | | | |

*(continued)*

| **[TG(19:1_18:0_18:1)+NH_4_]^+^** | M919T1066 | 918.84844 | 918.848415469999 | -0.6 | C5H9(69.070355):MS2,C5H11(71.085984):MS2,C6H9(81.070326):MS2,C6H11(83.085988):MS2,C6H13(85.101612):MS2,C7H9(93.070071):MS2,C7H11(95.085824):MS2,C7H13(97.101434):MS2,C8H11(107.085542):MS2,C8H13(109.101342):MS2,C8H15(111.116702):MS2,C9H13(121.101214):MS2,C9H15(123.116826):MS2,C10H15(135.116899):MS2,C10H17(137.132673):MS2,C11H17(149.132733):MS2,FA(18:1)-OH(265.252161):MS2,FA(19:1)-OH(279.26764):MS2,NL[FA(19:1)-H+NH4](605.549317):MS2,NL[FA(18:0)-H+NH4](617.550518):MS2,NL[FA(18:1)-H+NH4](619.565857):MS2 |
| --- | --- | --- | --- | --- | --- |
| §Unconfirmed in ESI negative mode | | | | | |
| **[TG(19:0_18:1_18:1)+Na]^+^** | M924T1066 | 923.80341 | 923.803811469999 | -1.0 | C6H9(81.070346):MS2,C6H11(83.086041):MS2,C6H13(85.101671):MS2,C7H11(95.085862):MS2,C7H13(97.101444):MS2,C8H13(109.101423):MS2,C8H15(111.116976):MS2,C9H13(121.101457):MS2,C9H15(123.117067):MS2,C10H15(135.116892):MS2,FA(18:1)-OH(265.252952):MS2,NL[FA(19:0)-H+Na](603.537095):MS2,NL[FA(18:1)-H+Na](619.566554):MS2,NL[FA(19:0)](625.517251):MS2,NL[FA(18:1)](641.549928):MS2,M+Na(923.804176):MS2 |
| §Unconfirmed in ESI negative mode | | | | | |
| † Lipid relevant from both ESI+ and ESI- datasets  § Lipid class only detected in ESI positive mode [7], cannot be confirmed by ESI negative mode | | | | | |

**Supplementary Table 2**. Statistical results from the Lipidyzer platform. Only the significant lipids are presented. Light green: *p*-value ≤ 0.06; Green: *p*-value ≤ 0.05; Yellow: *p*-value ≤ 0.01

| **Lipid species** | ***p-*value D3** | ***p-*value D18** | ***p-*value D23** |
| --- | --- | --- | --- |
| **CE** | | | |
| CE(14:0) | 0.42 | 0.03 | 0.22 |
| CE(14:1) | 1.00 | 0.13 | 0.02 |
| CE(15:0) | 0.84 | 0.03 | 0.15 |
| CE(16:0) | 0.69 | 0.03 | 0.10 |
| CE(16:1) | 0.84 | 0.03 | 0.10 |
| CE(17:0) | 1.00 | 0.03 | 0.10 |
| CE(18:0) | 0.22 | 0.03 | 0.31 |
| CE(18:1) | 0.55 | 0.03 | 0.10 |
| CE(18:2) | 0.42 | 0.03 | 0.10 |
| CE(18:3) | 0.55 | 0.03 | 0.15 |
| CE(18:4) | 0.69 | 0.03 | 0.03 |
| CE(20:0) | 1.00 | 0.03 | 0.55 |
| CE(20:1) | 0.84 | 0.03 | 0.55 |
| CE(20:2) | 0.15 | 0.03 | 0.22 |
| CE(20:4) | 0.42 | 0.03 | 0.15 |
| CE(22:0) | 0.22 | 0.03 | 0.06 |
| CE(22:1) | 0.84 | 0.03 | 0.10 |
| CE(22:2) | 0.55 | 0.03 | 0.15 |
| CE(22:4) | 0.84 | 0.03 | 0.31 |
| CE(22:5) | 0.31 | 0.03 | 0.22 |
| CE(24:0) | 1.00 | 0.03 | 0.10 |
| CE(24:1) | 0.42 | 0.03 | 0.31 |
| **CER** | | | |
| CER(24:0) | 0.42 | 0.03 | 0.22 |
| **DAG** | | | |
| DAG(16:0_18:2) | 0.55 | 0.49 | 0.01 |
| DAG(16:0_20:4) | 0.89 | 0.03 | 0.70 |
| DAG(16:1_18:1) | 0.10 | 0.69 | 0.02 |
| DAG(16:1_18:2) | 0.69 | 0.20 | 0.02 |
| DAG(18:0_18:2) | 1.00 | 0.49 | 0.01 |
| DAG(18:1_18:1) | 0.69 | 0.34 | 0.02 |
| DAG(18:1_18:2) | 0.42 | 0.34 | 0.02 |
| DAG(18:1_20:1) | 0.57 | 0.86 | 0.01 |
| DAG(18:1_20:4) | 0.69 | 0.11 | 0.02 |
| DAG(18:2_18:3) | 0.86 | 0.40 | 0.04 |
| DAG(18:2_20:4) | 0.55 | 0.34 | 0.01 |
| **DCER** | | | |
| DCER(24:1) | 0.69 | 0.03 | 0.06 |
| **FFA** | | | |
| FFA(20:5) | 0.22 | 0.03 | 0.03 |
| **HCER** | | | |
| HCER(16:0) | 0.42 | 0.03 | 0.42 |
| HCER(24:0) | 0.42 | 0.03 | 0.55 |
| HCER(24:1) | 0.03 | 0.03 | 0.55 |

Supplementary Table 2 (continued).

| **Lipid species** | ***p-*value D3** | ***p-*value D18** | ***p-*value D23** |
| --- | --- | --- | --- |
| **LCER** | | | |
| LCER(16:0) | 0.55 | 0.03 | 0.31 |
| LCER(18:0) | 0.84 | 0.03 | 0.19 |
| LCER(20:1) | 0.92 | 0.06 | 0.03 |
| **LPE** | | | |
| LPE(18:0) | 1.00 | 0.03 | 0.55 |
| **PC** | | | |
| PC(14:0_18:1) | 0.42 | 0.03 | 0.06 |
| PC(14:0_18:2) | 0.84 | 0.03 | 0.15 |
| PC(15:0_18:1) | 0.41 | 0.03 | 0.22 |
| PC(15:0_18:2) | 0.56 | 0.03 | 0.06 |
| PC(16:0_14:0) | 0.73 | 0.03 | 0.11 |
| PC(16:0_16:0) | 0.22 | 0.03 | 0.31 |
| PC(16:0_16:1) | 1.00 | 0.03 | 0.01 |
| PC(16:0_18:0) | 0.06 | 0.03 | 0.84 |
| PC(16:0_18:1) | 1.00 | 0.03 | 0.01 |
| PC(16:0_18:2) | 0.42 | 0.03 | 0.10 |
| PC(16:0_18:3) | 0.42 | 0.03 | 0.06 |
| PC(16:0_20:1) | 0.29 | 0.03 | 0.22 |
| PC(16:0_20:2) | 0.69 | 0.03 | 0.31 |
| PC(16:0_20:4) | 0.84 | 0.03 | 0.03 |
| PC(16:0_22:5) | 0.69 | 0.03 | 0.15 |
| PC(17:0_18:1) | 0.69 | 0.03 | 0.03 |
| PC(17:0_18:2) | 1.00 | 0.03 | 0.15 |
| PC(17:0_20:4) | 0.22 | 0.03 | 0.06 |
| PC(18:0_18:0) | 0.31 | 0.11 | 0.02 |
| PC(18:0_18:1) | 0.84 | 0.20 | 0.03 |
| PC(18:1_16:1) | 0.69 | 0.03 | 0.03 |
| PC(18:1_18:1) | 0.69 | 0.06 | 0.01 |
| PC(18:1_18:3) | 0.06 | 0.03 | 0.06 |
| PC(18:1_20:4) | 1.00 | 0.03 | 0.03 |
| PC(18:2_16:1) | 0.55 | 0.03 | 0.10 |
| PC(18:2_20:4) | 0.22 | 0.03 | 0.06 |
| **PE** | | | |
| PE(16:0_18:2) | 0.55 | 0.11 | 0.02 |
| PE(16:0_20:4) | 0.69 | 0.03 | 0.02 |
| PE(18:0_18:3) | 0.57 | 0.86 | 0.04 |
| PE(18:1_18:2) | 0.84 | 0.11 | 0.02 |
| PE(18:2_16:1) | 0.40 | 0.03 | 0.11 |
| PE(O-18:0_18:1) | 0.02 | 0.03 | 0.15 |
| PE(O-18:0_20:4) | 0.31 | 0.03 | 0.01 |
| PE(P-16:0_20:4) | 0.22 | 0.11 | 0.02 |
| PE(P-18:0_18:1) | 1.00 | 0.03 | 0.22 |
| PE(P-18:0_18:2) | 0.84 | 0.03 | 0.06 |
| PE(P-18:1_18:2) | 1.00 | 0.03 | 0.22 |
| PE(P-18:1_20:4) | 0.69 | 0.11 | 0.03 |

Supplementary Table 2 (continued).

| **Lipid species** | ***p*-value D3** | ***p*-value D18** | ***p*-value D23** |
| --- | --- | --- | --- |
| **SM** | | | |
| SM(16:0) | 0.10 | 0.03 | 0.84 |
| SM(22:1) | 0.42 | 0.03 | 0.55 |
| SM(24:0) | 0.69 | 0.03 | 0.22 |
| SM(24:1) | 0.42 | 0.03 | 0.15 |
| SM(26:0) | 0.84 | 0.03 | 0.06 |
| **TG** | | | |
| TG42:1-FA14:0 | 0.06 | 1.00 | 0.02 |
| TG44:0-FA12:0 | 0.22 | 0.11 | 0.03 |
| TG44:0-FA14:0 | 0.69 | 0.03 | 0.42 |
| TG44:1-FA16:0 | 1.00 | 0.03 | 0.04 |
| TG44:1-FA18:1 | 0.56 | 0.03 | 0.04 |
| TG44:2-FA16:0 | 0.73 | 0.03 | 0.03 |
| TG44:2-FA18:2 | 0.56 | 0.03 | 0.03 |
| TG45:0-FA15:0 | 0.73 | 0.03 | 0.10 |
| TG45:0-FA16:0 | 0.84 | 0.03 | 0.22 |
| TG45:1-FA18:1 | 1.00 | 0.50 | 0.01 |
| TG46:1-FA18:1 | 0.55 | 0.20 | 0.02 |
| TG46:2-FA14:0 | 0.22 | 0.06 | 0.01 |
| TG46:2-FA18:1 | 1.00 | 0.03 | 0.02 |
| TG46:3-FA12:0 | 0.73 | 0.11 | 0.03 |
| TG46:3-FA14:1 | 0.42 | 0.03 | 0.03 |
| TG46:3-FA18:1 | 0.73 | 0.03 | 0.01 |
| TG46:3-FA18:2 | 0.42 | 0.03 | 0.15 |
| TG47:0-FA14:0 | 1.00 | 0.03 | 0.11 |
| TG47:1-FA16:0 | 1.00 | 0.03 | 0.31 |
| TG47:1-FA18:1 | 0.42 | 0.06 | 0.03 |
| TG47:2-FA14:0 | 0.84 | 0.11 | 0.02 |
| TG47:2-FA15:0 | 0.69 | 0.03 | 0.01 |
| TG47:2-FA16:1 | 0.31 | 0.03 | 0.31 |
| TG47:2-FA18:1 | 0.84 | 0.03 | 0.03 |
| TG47:2-FA18:2 | 0.56 | 0.03 | 0.15 |
| TG48:0-FA16:0 | 1.00 | 0.11 | 0.03 |
| TG48:1-FA12:0 | 1.00 | 0.23 | 0.03 |
| TG48:1-FA18:0 | 1.00 | 0.20 | 0.01 |
| TG48:2-FA12:0 | 0.42 | 0.11 | 0.02 |
| TG48:2-FA14:1 | 0.84 | 0.03 | 0.03 |
| TG48:2-FA16:0 | 0.84 | 0.06 | 0.02 |
| TG48:2-FA18:1 | 1.00 | 0.11 | 0.03 |
| TG48:3-FA16:0 | 0.55 | 0.06 | 0.03 |
| TG48:3-FA16:1 | 1.00 | 0.06 | 0.02 |
| TG48:3-FA18:3 | 0.42 | 0.11 | 0.02 |
| TG48:4-FA14:0 | 0.49 | 0.23 | 0.02 |
| TG48:4-FA14:1 | 1.00 | 0.67 | 0.04 |
| TG48:4-FA16:0 | 1.00 | 0.11 | 0.02 |
| TG49:1-FA14:0 | 0.31 | 0.03 | 0.10 |

Supplementary Table 2 (continued).

| **Lipid species** | ***p-*value D3** | ***p-*value D18** | ***p-*value D23** |
| --- | --- | --- | --- |
| **TG (continued)** | | | |
| TG49:1-FA16:1 | 0.69 | 0.03 | 0.15 |
| TG49:2-FA14:0 | 0.42 | 0.03 | 0.15 |
| TG49:2-FA17:0 | 0.42 | 0.03 | 0.01 |
| TG50:1-FA16:0 | 1.00 | 0.20 | 0.03 |
| TG50:2-FA14:0 | 0.84 | 0.11 | 0.03 |
| TG50:2-FA14:1 | 0.57 | 0.06 | 0.01 |
| TG50:2-FA18:0 | 0.69 | 0.11 | 0.02 |
| TG50:3-FA14:1 | 1.00 | 0.06 | 0.03 |
| TG50:3-FA18:0 | 0.55 | 0.06 | 0.02 |
| TG50:4-FA14:1 | 0.42 | 0.03 | 0.06 |
| TG50:4-FA16:0 | 0.55 | 0.34 | 0.01 |
| TG50:4-FA18:1 | 0.55 | 0.11 | 0.01 |
| TG50:4-FA20:4 | 0.86 | 0.20 | 0.03 |
| TG50:5-FA16:0 | 1.00 | 0.06 | 0.01 |
| TG50:5-FA16:1 | 1.00 | 0.34 | 0.01 |
| TG50:5-FA18:1 | 0.55 | 0.06 | 0.03 |
| TG50:5-FA18:3 | 0.31 | 0.06 | 0.01 |
| TG51:2-FA16:0 | 0.55 | 0.06 | 0.03 |
| TG51:2-FA16:1 | 0.55 | 0.03 | 0.06 |
| TG51:2-FA17:0 | 0.84 | 0.03 | 0.10 |
| TG51:2-FA18:2 | 0.84 | 0.03 | 0.10 |
| TG51:3-FA16:1 | 0.31 | 0.03 | 0.06 |
| TG51:3-FA17:0 | 0.55 | 0.03 | 0.06 |
| TG51:4-FA16:1 | 0.31 | 0.03 | 0.01 |
| TG52:0-FA20:0 | 1.00 | 0.06 | 0.02 |
| TG52:1-FA20:0 | 1.00 | 0.03 | 0.05 |
| TG52:2-FA14:0 | 0.55 | 0.03 | 0.01 |
| TG52:2-FA20:0 | 0.69 | 0.11 | 0.01 |
| TG52:2-FA20:1 | 0.84 | 0.03 | 0.01 |
| TG52:3-FA14:0 | 1.00 | 0.06 | 0.01 |
| TG52:3-FA16:1 | 1.00 | 0.11 | 0.02 |
| TG52:3-FA20:1 | 0.84 | 0.06 | 0.01 |
| TG52:3-FA20:2 | 0.55 | 0.20 | 0.03 |
| TG52:4-FA18:0 | 0.69 | 0.34 | 0.03 |
| TG52:4-FA20:0 | 0.42 | 0.06 | 0.03 |
| TG52:4-FA20:2 | 0.40 | 0.06 | 0.01 |
| TG52:4-FA22:4 | 0.80 | 0.67 | 0.04 |
| TG52:5-FA20:3 | 0.84 | 0.03 | 0.01 |
| TG52:6-FA18:1 | 0.42 | 0.34 | 0.02 |
| TG52:6-FA20:4 | 0.69 | 0.06 | 0.01 |
| TG52:6-FA20:5 | 1.00 | 0.03 | 0.06 |
| TG52:7-FA20:5 | 0.90 | 0.11 | 0.02 |
| TG52:8-FA16:1 | 0.55 | 0.03 | 0.10 |
| TG52:8-FA18:2 | 1.00 | 0.03 | 0.06 |

Supplementary Table 2 (continued).

| **Lipid species** | ***p-*value D3** | ***p-*value D18** | ***p-*value D23** |
| --- | --- | --- | --- |
| **TG (continued)** | | | |
| TG53:1-FA18:0 | 0.69 | 0.03 | 0.10 |
| TG53:3-FA16:0 | 0.55 | 0.11 | 0.03 |
| TG53:3-FA17:0 | 1.00 | 0.03 | 0.10 |
| TG53:3-FA18:2 | 1.00 | 0.03 | 0.06 |
| TG53:4-FA16:0 | 0.42 | 0.06 | 0.01 |
| TG53:4-FA17:0 | 0.69 | 0.03 | 0.10 |
| TG53:4-FA18:2 | 1.00 | 0.03 | 0.06 |
| TG53:4-FA18:3 | 1.00 | 0.03 | 0.06 |
| TG53:4-FA20:4 | 1.00 | 0.06 | 0.01 |
| TG53:5-FA20:4 | 0.69 | 0.03 | 0.10 |
| TG54:0-FA16:0 | 0.84 | 0.03 | 0.01 |
| TG54:1-FA16:0 | 0.84 | 0.06 | 0.03 |
| TG54:1-FA20:0 | 0.84 | 0.06 | 0.01 |
| TG54:1-FA20:1 | 0.84 | 0.11 | 0.03 |
| TG54:2-FA16:0 | 0.69 | 0.06 | 0.02 |
| TG54:2-FA18:0 | 0.84 | 0.06 | 0.02 |
| TG54:2-FA20:0 | 0.55 | 0.06 | 0.01 |
| TG54:2-FA20:1 | 0.69 | 0.11 | 0.02 |
| TG54:3-FA16:0 | 0.42 | 0.06 | 0.01 |
| TG54:3-FA16:1 | 1.00 | 0.11 | 0.01 |
| TG54:3-FA18:1 | 0.69 | 0.11 | 0.01 |
| TG54:3-FA20:1 | 0.42 | 0.06 | 0.01 |
| TG54:4-FA16:1 | 0.55 | 0.06 | 0.01 |
| TG54:5-FA20:5 | 0.73 | 0.06 | 0.03 |
| TG54:6-FA22:6 | 0.31 | 0.06 | 0.01 |
| TG54:7-FA22:5 | 0.42 | 0.03 | 0.02 |
| TG54:7-FA22:6 | 0.84 | 0.20 | 0.01 |
| TG54:8-FA20:4 | 0.90 | 0.20 | 0.02 |
| TG54:8-FA20:5 | 0.84 | 0.03 | 0.10 |
| TG54:8-FA22:6 | 1.00 | 0.03 | 0.01 |
| TG55:1-FA18:1 | 0.42 | 0.03 | 0.10 |
| TG55:2-FA18:1 | 1.00 | 0.06 | 0.02 |
| TG55:3-FA18:1 | 0.55 | 0.03 | 0.02 |
| TG55:4-FA18:1 | 0.22 | 0.03 | 0.03 |
| TG55:5-FA18:1 | 0.84 | 0.06 | 0.01 |
| TG55:5-FA20:4 | 1.00 | 0.03 | 0.02 |
| TG55:7-FA15:0 | 1.00 | 0.03 | 0.06 |
| TG56:10-FA18:2 | 1.00 | 0.11 | 0.03 |
| TG56:1-FA16:0 | 0.31 | 0.03 | 0.01 |
| TG56:2-FA16:0 | 0.84 | 0.06 | 0.03 |
| TG56:2-FA18:0 | 0.84 | 0.03 | 0.02 |
| TG56:2-FA20:0 | 0.55 | 0.03 | 0.01 |
| TG56:2-FA20:1 | 0.69 | 0.03 | 0.02 |
| TG56:3-FA18:0 | 0.42 | 0.06 | 0.01 |
| TG56:3-FA18:1 | 0.22 | 0.06 | 0.02 |

Supplementary Table 2 (continued).

| **Lipid species** | ***p-*value D3** | ***p-*value D18** | ***p-*value D23** |
| --- | --- | --- | --- |
| **TG (continued)** | | | |
| TG56:3-FA20:0 | 0.42 | 0.06 | 0.01 |
| TG56:3-FA20:1 | 0.31 | 0.06 | 0.01 |
| TG56:3-FA20:2 | 0.31 | 0.06 | 0.03 |
| TG56:4-FA18:0 | 0.22 | 0.11 | 0.02 |
| TG56:4-FA18:1 | 0.22 | 0.06 | 0.01 |
| TG56:4-FA18:2 | 0.31 | 0.06 | 0.03 |
| TG56:4-FA20:1 | 0.42 | 0.06 | 0.03 |
| TG56:4-FA20:2 | 0.31 | 0.11 | 0.03 |
| TG56:4-FA20:3 | 0.42 | 0.11 | 0.02 |
| TG56:5-FA18:2 | 0.31 | 0.06 | 0.03 |
| TG56:6-FA18:3 | 0.42 | 0.11 | 0.01 |
| TG56:6-FA22:6 | 0.55 | 0.06 | 0.01 |
| TG56:7-FA18:0 | 0.55 | 0.03 | 0.10 |
| TG56:7-FA20:5 | 0.42 | 0.03 | 0.22 |
| TG56:8-FA18:1 | 0.42 | 0.03 | 0.10 |
| TG56:8-FA20:5 | 0.42 | 0.03 | 0.15 |
| TG56:9-FA20:5 | 0.69 | 0.03 | 0.06 |
| TG56:9-FA22:6 | 0.69 | 0.06 | 0.03 |
| TG57:2-FA18:1 | 0.42 | 0.06 | 0.03 |
| TG58:2-FA18:1 | 1.00 | 0.11 | 0.02 |
| TG58:3-FA18:1 | 0.42 | 0.06 | 0.03 |
| TG58:6-FA16:0 | 1.00 | 0.03 | 0.01 |
| TG58:7-FA16:0 | 1.00 | 0.03 | 0.10 |
| TG58:7-FA22:6 | 0.55 | 0.06 | 0.01 |
